# Structural basis for Rad54- and Hed1-mediated regulation of Rad51 during the transition from mitotic to meiotic recombination

**DOI:** 10.1101/2025.03.26.645561

**Authors:** Yeonoh Shin, Michael T. Petassi, Aidan M. Jessop, Stefan Y. Kim, Razvan Matei, Katherine Morse, Vivek B. Raina, Upasana Roy, Eric C. Greene

## Abstract

Rad51 catalyzes the DNA pairing reactions that take place during homologous recombination (HR), and HR must be tightly regulated to ensure physiologically appropriate outcomes. Rad54 is an ATP-dependent DNA motor protein that stimulates Rad51 activity during mitosis. In meiosis Rad51 is downregulated by the protein Hed1, which blocks Rad54 binding to Rad51, and allows Dmc1 to function as the active recombinase. We currently have a poor understanding of the regulatory interplay between Rad54, Hed1, Rad51 and Dmc1. Here, we identify a conserved Rad51 interaction motif within Rad54, and we solve a CryoEM structure of this motif bound to Rad51. We also identify a distinct Rad51 interaction motif within Hed1 and solve its structure bound to Rad51. These structures explain how Rad54 engages Rad51 to promote recombination between sister chromatids during mitosis and how Rad51 is downregulated by Hed1 upon entry into meiosis such that its meiosis-specific homolog Dmc1 can promote recombination between homologous chromosomes.

## INTRODUCTION

Homologous recombination (HR) is a universally conserved pathway used to repair double strand breaks (DSB), repair stalled and collapsed replication forks, and promote proper chromosome segregation and enhance genetic diversity during meiosis (Neale and Keeney, 2006; San Filippo et al., 2008; Symington et al., 2014). HR is essential for maintaining genome integrity and defects in HR are broadly associated with cancers and cancer-prone syndromes (Bonilla et al., 2020; Matos-Rodrigues et al., 2021).

The highly conserved Rad51/RecA family of proteins plays a central role in catalyzing the DNA pairing reactions that take place during HR (Bianco et al., 1998; Bonilla et al., 2020; Kowalczykowski, 2015). Rad51 is an ATP-dependent DNA-binding protein that forms extended right-handed helical filaments on the 3’ single stranded DNA (ssDNA) overhangs that are formed the processed ends of DSBs (Chen et al., 2008; Conway et al., 2004; Kowalczykowski, 2015; Morrical, 2015; San Filippo et al., 2008; Sheridan et al., 2008; Symington et al., 2014). The resulting Rad51-ssDNA nucleoprotein filament is also referred to as the presynaptic complex (Heyer et al., 2010; San Filippo et al., 2008; Symington et al., 2014). The Rad51 presynaptic complex is responsible for identifying a homologous double stranded DNA (dsDNA) sequence that can be used as a template to guide repair of the damaged DNA, and it catalyzes a strand invasion reaction where it pairs the bound ssDNA with the complementary strand of the homologous dsDNA, resulting in displacement of the non–complementary DNA strand. The resulting D-loop intermediate can be processed through several different pathways, leading to repair of the damaged DNA (Heyer et al., 2010; Mehta and Haber, 2014; Paques and Haber, 1999; San Filippo et al., 2008; Symington et al., 2014).

Numerous accessory proteins are required to both positively and negatively regulate Rad51 during HR. Rad54 is a member of the Swi/Snf family of ATP-dependent dsDNA translocases and is also one of the most highly conserved HR regulatory proteins found in eukaryotes (Ceballos and Heyer, 2011b; Heyer et al., 2006; Kowalczykowski, 2015; Mazin et al., 2010b). *RAD54* deletion imparts sensitivity to DNA damaging agents (Petukhova et al., 1999; Wesoly et al., 2006), causes defects in strand invasion (Renkawitz et al., 2013; Sugawara et al., 2003), leads to the accumulation of toxic HR intermediates (Shah et al., 2010a), and *rad54* missense mutations have been linked to human cancers (Matsuda et al., 1999; Sridalla et al., 2024). Rad54 binds tightly to the Rad51 presynaptic complex, drives the ATP-hydrolysis-dependent movement of the presynaptic complex along dsDNA during the search for sequence homology, and it greatly increases the efficiency of Rad51-mediated strand invasion (Crickard et al., 2020b; Jiang et al., 1996; Kowalczykowski, 2015; Mazin et al., 2000; Petukhova et al., 1998; Raschle et al., 2004; Van Komen et al., 2000). In addition to its functions in promoting the homology search and strand invasion, Rad54 has been implicated in the stabilization of the Rad51 presynaptic complex (Mazin et al., 2003a; Meir et al., 2022), DNA branch migration (Bugreev et al., 2006; Goyal et al., 2018; Rossi and Mazin, 2008), nucleosome remodeling (Alexeev et al., 2003; Alexiadis and Kadonaga, 2002; Jaskelioff et al., 2003), and the removal of Rad51 from dsDNA (Ceballos and Heyer, 2011b; Symington and Heyer, 2006; Wright and Heyer, 2014).

The transition from mitotic to meiotic recombination involves a change in repair template preference: in mitosis, recombination occurs preferentially between sister chromatids, while in meiosis, it takes place more frequently between homologous chromosomes (Arter and Keeney, 2024; Brown and Bishop, 2014; Lao and Hunter, 2010). Meiotic recombination also involves the selective downregulation of Rad51 activity, allowing the meiosis-specific recombinase Dmc1 to catalyze strand exchange (Brown and Bishop, 2014; Cloud et al., 2012; Da Ines et al., 2013; Hinch et al., 2020; Neale and Keeney, 2006; Petiot et al., 2024; Singh et al., 2017). Therefore, Dmc1 is the active recombinase during meiosis, with Rad51 acting as an accessory factor to facilitate Dmc1 filament assembly (Brown and Bishop, 2014; Cloud et al., 2012). Dmc1-mediated repair of the programmed DSBs that are generated by Spo11 during meiosis creates physical linkages between homologous chromosomes called chiasma, which are essential to ensure proper chromosome segregation (Brown and Bishop, 2014; Lam and Keeney, 2014; Lao and Hunter, 2010; Zickler and Kleckner, 2015). In *S. cerevisiae*, Rad51 inhibition is achieved through two meiosis-specific regulatory proteins, Mek1 and Hed1 (Brown and Bishop, 2014; Niu et al., 2009). Mek1 is a kinase that phosphorylates Rad54 to weaken its interaction with Rad51 (Niu et al., 2009) and Hed1 is a regulatory protein that binds to Rad51 and blocks the association of Rad54 (Busygina et al., 2012; Busygina et al., 2008). Rad54 is a required co-factor for Rad51 strand invasion activity, therefore, Hed1 binding downregulates the activity of Rad51 in meiosis by preventing interactions between Rad51 and Rad54. Interestingly, even though *S. cerevisiae* Dmc1 and Rad51 both interact with Rad54, the negative regulatory function of Hed1 is highly specific for just Rad51 (Busygina et al., 2008; Crickard et al., 2018; Tsubouchi and Roeder, 2006).

We currently have a limited understanding of the mechanistic interplay that underlies the regulation of Rad51 by cofactors such as Rad54 and Hed1 largely because there are no high-resolution structures of Rad51 presynaptic complexes bound to these cofactors. Here, we use a combination of bioinformatics and AlphaFold3 modeling to identify a broadly conserved peptide within the unstructured N-terminal domain of Rad54 that is predicted to interact with Rad51. We then solve the CryoEM structure of this Rad51 interaction motif bound to the Rad51-ssDNA filament and use a combination of deep mutagenesis and site-directed mutagenesis to validate important amino acid residue contacts, providing structural insights into Rad54 protein-protein interactions with the Rad51 presynaptic complex. We use a similar approach to predict how Hed1 binds to Rad51, we then solve the CryoEM structure of the Rad51 interaction motif from Hed1 bound to a Rad51-ssDNA filament and we validate important amino acid residues using mutagenesis. Our data demonstrate that the Hed1 binding pocket physically overlaps with the Rad54 binding site, explaining how Hed1 acts as a potent negative regulatory factor of Rad51 strand exchange activity during meiosis. Notably, although their binding sites overlap, the binding mechanisms of Rad54 and Hed1 are completely different from one another. These distinct binding mechanisms help to explain how Rad54 can interact with both Rad51 and Dmc1, whereas Hed1 selectively binds to just Rad51 and does not bind to Dmc1, even though Rad51 and Dmc1 share 45% sequence identity, thus allowing Dmc1 to act as the primary recombinase for promoting recombination between homologous chromosomes during meiosis.

## RESULTS

### Predicted interaction between Rad54 and Rad51

Crystal structures of the core Snf2 motor domain from *Danio rerio* Rad54 and a Rad54 homolog from *Sulfolobus solfataricus* have been reported (Dürr et al., 2005; Thomä et al., 2005). However, there are currently no structures showing how Rad54 interacts with the Rad51 presynaptic complex. Initial efforts at obtaining CryoEM structures of *S. cerevisiae* Rad51 bound to full-length *S. cerevisiae* Rad54 (898 aa) were not successful, therefore we sought to obtain a structure of a smaller complex that would still enable us to define how Rad54 interacts with the Rad51-ssDNA presynaptic filament. For this, we used a combination of bioinformatics and AlphaFold3 modeling to predict which region of Rad54 might be involved in direct contacts with Rad51 (Figure 1). Rad54 is comprised of a largely unstructured N-terminal domain (NTD) that interacts with Rad51 and a C-terminal domain (CTD) containing an Snf2 motor domain (Figure 1A)(Ceballos and Heyer, 2011a; Crickard and Greene, 2019; Crickard et al., 2020a; Mazin et al., 2010a). Bioinformatic analysis revealed the existence of a conserved region (amino acids 101 to 144, Figure 1B) within the Rad54 NTD that also contains an FxxP motif conserved across eukaryotes (Figure 1C). As anticipated, AlphaFold3 predictions suggest that the Rad54 NTD is largely unstructured (Figure S1A-S1B), which is consistent with previous studies (Raschle et al., 2004). However, AlphaFold3 predictions also suggested that a portion of the NTD containing the FxxP motif becomes more highly structured in the presence of Rad51 and can interact with the Rad51-ssDNA nucleoprotein filament (Figure 1D-1E & Figure S1C-S1D). Given these considerations, we tentatively defined *S. cerevisiae* Rad54 amino acid residues T101 to L144 as a potential Rad51 interaction motif.

**Figure 1.**
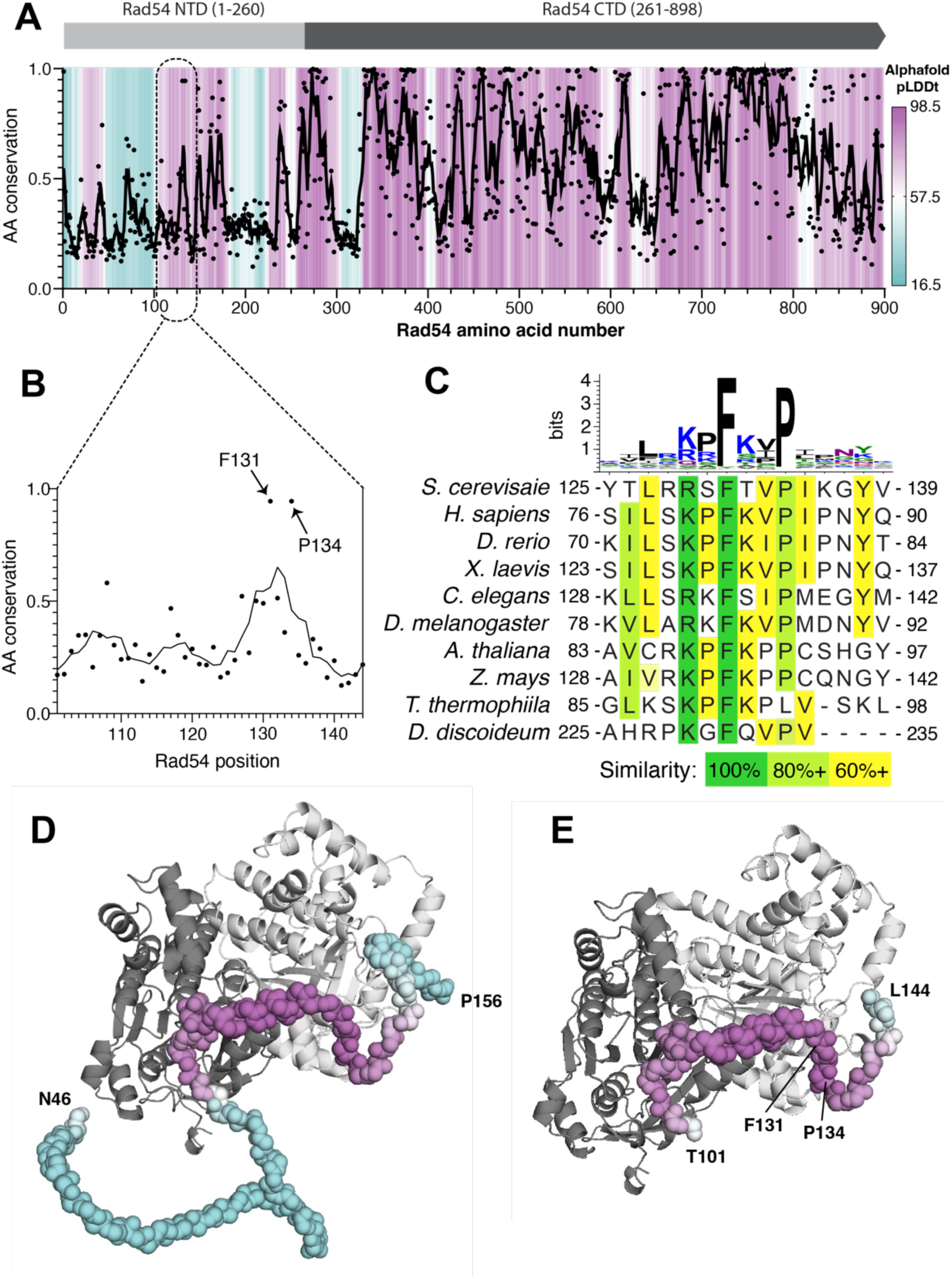
Predicted interaction interface between Rad51 and Rad54. **(A)** Plot of amino acid residue conservation and predicted AlphaFold3 structure (pLDDt; predicted local difference distance test) versus residue number for *S. cerevisiae* Rad54. Individual amino acid positions are indicated as black dots with a 5-amino acid moving average plotted as a black line. pLDDt is colored from cyan to purple as indicated by the scale bar. **(B)** Residue conservation in the region of the N-terminal domain FxxP motif. **(C)** Sequence logo and sequence alignments for the region encompassing the FxxP motif. **(D)** Predicted AlphaFold3 structure for a Rad54 residues N46 to P156 or **(E)** Rad54 residues T101-L144 bound to two adjacent Rad51 monomers from within a Rad51 filament.

### Structure of the Rad54-bound presynaptic complex

We obtained a 3.3 Å resolution CyroEM structure of an *S. cerevisiae* Rad51 filament bound to a 96-nuclotide ssDNA fragment in the presence of ATP along with a 44 amino acid peptide encompassing residues T101 to L144 from Rad54 (Figure 2, Figure S2, Figure S3A-B, & Table S1). The Rad54 peptide decorates the outside surface of the Rad51 filament, and it is located on the opposite side of the nucleoprotein filament relative to the bound strand of ssDNA (Figure 2A & 2B). Each Rad54 peptide interacts with a composite binding site formed by two adjacent Rad51 protomers within the nucleoprotein filament, corresponding to a buried surface area of 1651.4 Å^2^ (Figure 2A & 2C). The Rad54 peptide is comprised of a N-terminal loop (amino acid residues R103 to A110), followed by a short helix (∼1 turn; residues Q111 to D116), then another short helix (∼3 turns; residues P117 to T126), and ends with a loop region (residues L127 to K136) that contains the FxxP motif and is located near the C-terminus of the peptide. Rad54 residues R103 to T126 interact with the Rad51 protomer facing towards the 3’ end of the ssDNA (shown in cyan; Figure 2D-F) and residues R128-K136 interact with the 5’ facing Rad51 protomer (shown in green; Figure 2D-F). Close inspection of the structure reveals that the Rad54 peptide interacts with Rad51 primarily through a combination of hydrophobic and electrostatic contacts. There are two hydrophobic regions on the surface of the Rad51 filament that interact with the Rad54 peptide. Rad51 residues V140 to V145 form a hydrophobic pocket at the Rad51 protomer-protomer interface which interacts with leucine residues L113, L120, and L127 and isoleucine residues I123 from the Rad54 peptide (Figure 2D). Rad51 residues G209 to G213 form a second hydrophobic pocket which interacts with the FxxP motif of Rad54 (Figure 2D). Notably, these hydrophobic pockets are highly conserved in both the Rad51 and Dmc1 lineages of the Rad51/RecA family of proteins, suggesting that this may be a general mechanism for Rad54 binding to the surface of these two eukaryotic recombinases (*see Discussion*). In addition to these hydrophobic contacts, the basic residues K105, R112, R119, R128 and R129 from Rad54 form a network of electrostatic interactions with Rad51 residues E212, E156, D149, E271, and N246 (Figure 2E). For brevity, we will hereafter refer to the 44 amino acid residue peptide (T101 to L144) as the Rad51 interaction motif of Rad54.

**Figure 2.**
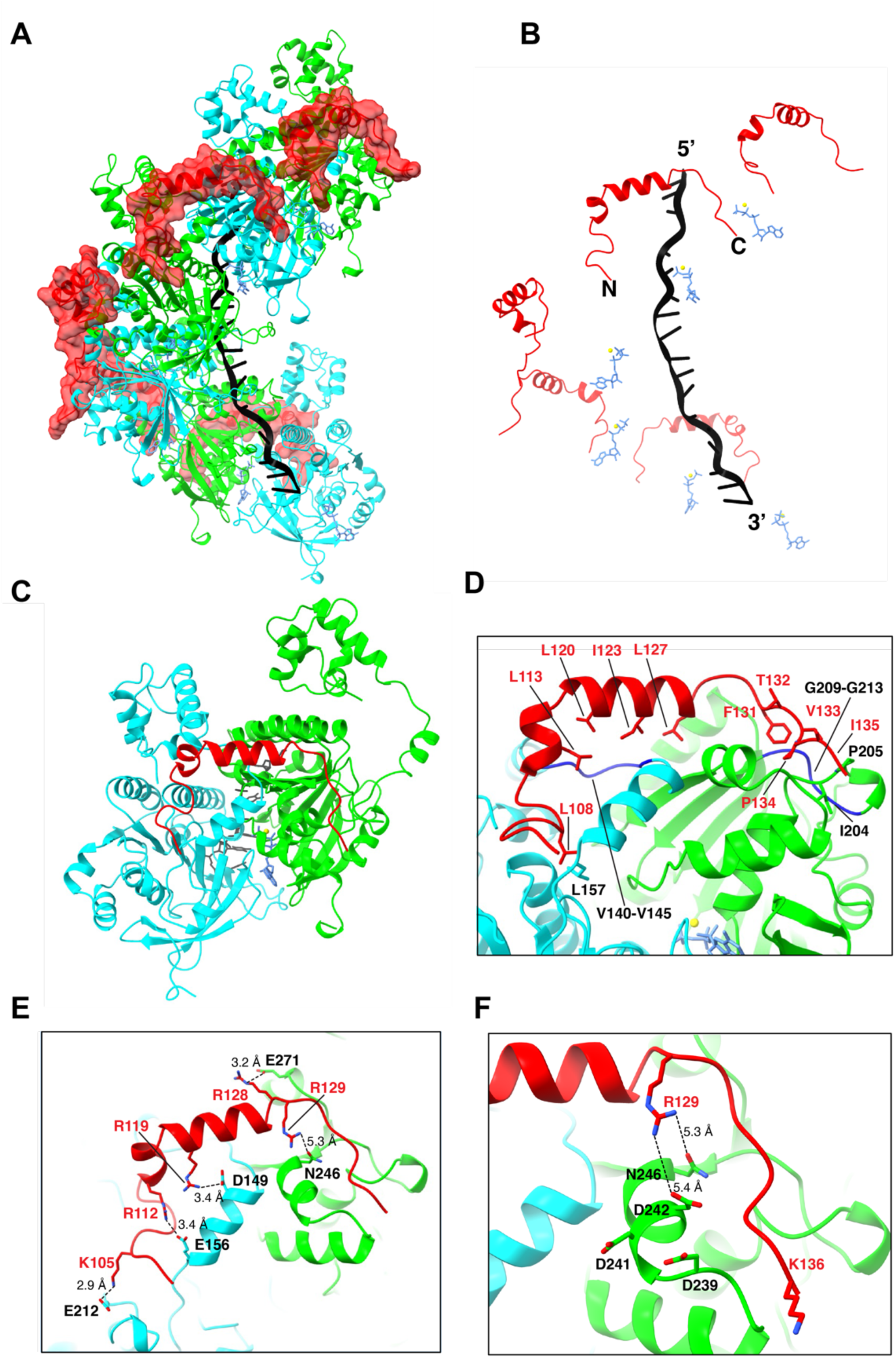
Structure of the Rad51 interaction motif of Rad54 bound to the Rad51-ssDNA filament. **(A)** CryoEM reconstruction of the Rad51-ssDNA filament bound by a peptide encompassing Rad54 residues T101 to L144. Rad51 subunits are shown in alternating colors (cyan and green) and the Rad54 peptide is shown in red. **(B)** Structure in which the Rad51 proteins have been removed to highlight the relative locations of the Rad54 peptide, Rad51-bound ssDNA and ATP molecules. **(C)** Structure showing that the Rad54 peptide is bound to two adjacent Rad51 subunits (the 5’ oriented subunit is shown in green and the 3’ oriented subunit is shown in cyan). **(D-F)** Details of the molecular interactions between Rad54 (shown in red) and the two adjacent Rad51 subunits.

### Analysis of the Rad51 conserved protruding acidic patch

It has been proposed that a conserved protruding acidic patch (PAP) on the surface of Rad51, comprised of residues D239, D241 and D242, was involved in interactions with Rad54 (Afshar et al., 2021). Surprisingly, these PAP residues do not appear to form obvious strong interactions with the Rad51 interaction motif from the Rad54 peptide in our structure (Figure 2F). Rad51 residue D241 faces away from the bound Rad54 peptide, and Rad51 residues D239 and D242 are too far from the nearest basic residues in Rad54 to form strong electrostatic interactions (Figure 2F). To test the potential contributions of these acidic Rad51 residues to interactions with Rad54 we tested *rad51* mutants in which the PAP aspartic acid residues were changed to alanine and asked whether cells expressing these mutants could survive when grown on media containing the DNA damaging agent methyl methane sulfonate (MMS). The *rad51* single mutants (D239A; D241A; and D242A) and *rad51* double mutants (D241A, D242A; D239A, D241A; and D239A, D242A) all exhibit wild-type or near wild-type phenotypes when grown on media containing the DNA damaging agent methyl methanesulfonate (MMS), whereas a *rad51* triple mutant (D239A, D241A, D242A) exhibits a null phenotype (Figure S3C). However, the triple mutant Rad51 protein still supports Rad54-dependent D-loop formation *in vitro* suggesting that the *in vivo* defect likely is not directly related to the disruption of Rad51 interactions with Rad54 (Figure S3D). We noted that the Rad51 triple mutant (D239A, D241A, D242A) precipitates during purification at salt concentrations lower than 500 mM NaCl, suggesting the possibility that in our genetic assays the three mutations may act to destabilize the protein rather than disrupting any specific interactions with Rad54. Alternatively, the triple mutant protein may be defective in cells due to disruption of interactions with other proteins such as Rad55-Rad57, Rad52, or both (Afshar et al., 2021). Taken together, our data suggest that the Rad51 protruding acid patch might not be directly involved in contacts between Rad51 and Rad54.

### Genetic analysis of the Rad51 interaction motif from Rad54

We used site directed mutagenesis to assess the importance of the interfacial contacts between Rad51 and Rad54 observed in the CryoEM structure. For this analysis, we used our CryoEM structure to guide the design of *rad54* mutants with various combinations of single or multiple point mutations at amino acid residues shown to make contacts with Rad51 (Figure 2D-F). These *rad54* mutants were cloned in a CEN vector and their functionality was assessed by determining whether cells expressing these mutants could survive when grown on media containing MMS (Figure S4). Interestingly, most single point mutants within the Rad51 interaction motif of Rad54 had little or no effect on cell survival (Figure S4A). The sole exceptions were L127A, L127D and F131A. Residue L127 makes hydrophobic contacts with a conserved region of Rad51 (Figure 2D) and residue F131 is part of the FxxP motif and phenylalanine was strongly preferred at this position (Figure 2D & Figure 3B-C). Notably, the P134A mutation, which is also part of the FxxP motif, was tolerated in the point mutant assays (*see below*; Figure S4).

**Figure 3.**
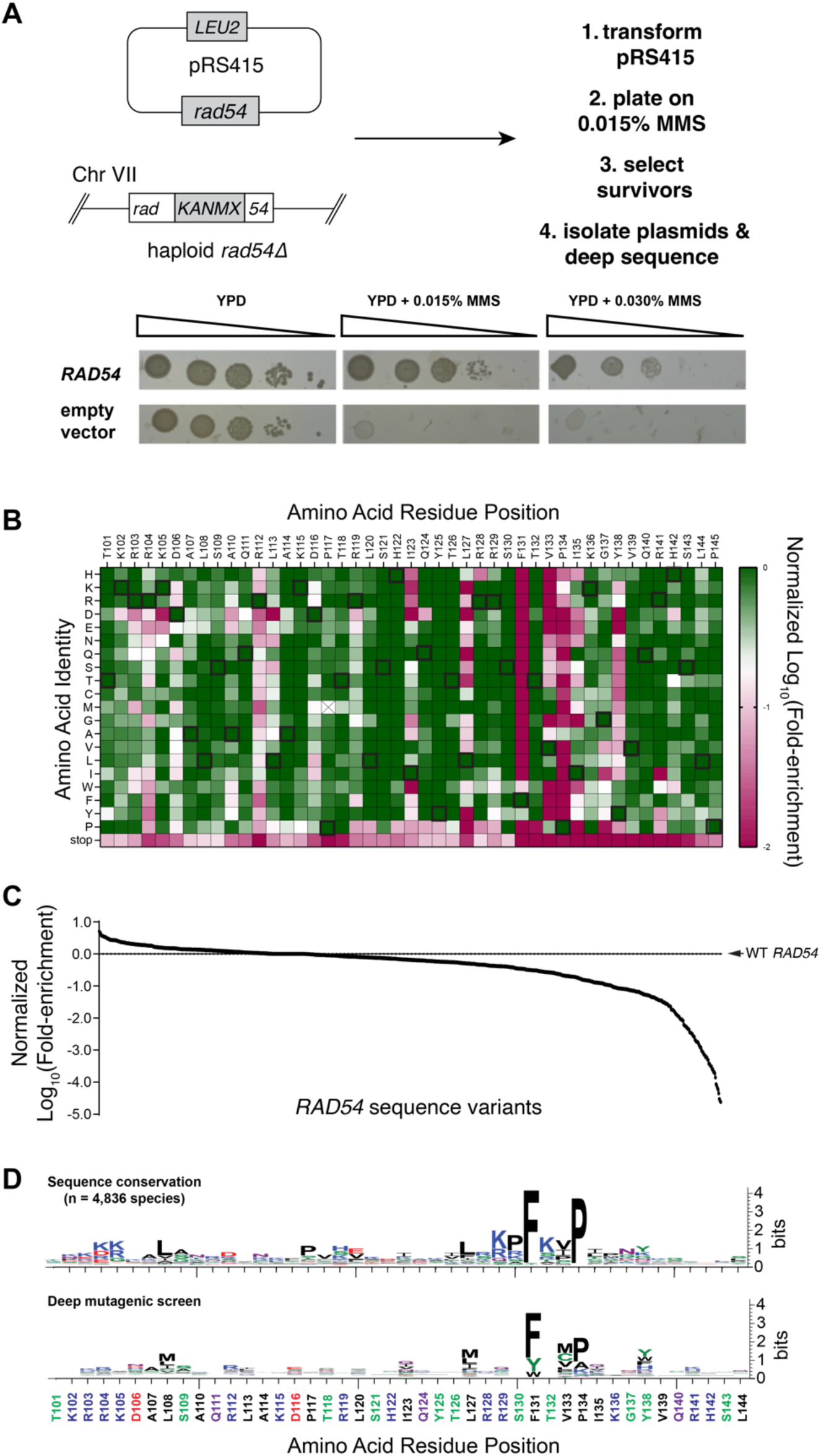
Functional landscape of the Rad54-Rad51 interface. **(A)** Strategy for deep mutagenic analysis of the Rad51 interaction motif from Rad54 and control spot assays on MMS-containing media for a *rad54Δ* strain complemented with either wild-type RAD54 or an empty vector. **(B)** Heat map showing Rad54 single mutant sequence variants that were recovered from the deep mutagenic screen. Data is normalized to the wild-type allele at each position and colored from green (corresponding to wild-type), to white (Log_10_(−0.7), corresponding to ∼5-fold depleted relative to wild-type), to red (Log_10_(−2), corresponding to 100-fold depleted relative to wild-type). The P117M allele, marked with an X, was underrepresented in the initial pool so it was omitted from further analysis. **(C)** Distribution of all data values for the deep mutagenic screen. **(D)** Sequence logos comparing the Rad54 sequence variation found in nature (top panel) with the results of the deep mutagenic screen (bottom panel).

In contrast to the individual point mutations, all *rad54* alleles bearing multiple mutations exhibited genetic defects relative to wild-type *RAD54* (Figure S4B). Four of these contain the L127A mutation, which alone likely accounts for the observed defects (Figure S4B). Additionally, other alleles with multiple mutations in interacting residues yielded significantly reduced growth on MMS-containing plates comparable to the F131A point mutant (Figure S4A-S4B). Examples of these highly defective mutants include: the *rad54* triple mutant K105D R112D R119D, all of which are located near the N-terminal end of the Rad51 interaction domain (Figure 2E); the *rad54* triple mutant R128D R129D K136D, all located near the C-terminal end of the Rad51 interaction domain (Figure 2E-F), and the *rad54* triple mutant R129A V133A I135A, which are also clustered near the C-terminal end of the Rad51 interaction motif (Figure 2D-2F).

### Deep mutagenic screen of the Rad51 interaction motif from Rad54

Given that most single point mutations with the Rad51 interaction domain of Rad54 were tolerated, we next conducted a deep mutagenic screen to help further define the functional significance of the amino acid residues within the Rad51 interaction motif of Rad54. For this assay, we generated libraries of Rad54 mutants in which each amino acid across the entire Rad51 interaction motif was mutated to all other possible amino acid residues (Figure 3A). The *rad54* mutant libraries were encoded within a CEN plasmid (*pRS415–ScRad54*) in which the codons from within the 44 amino acid residue Rad51 interaction motif (spanning Rad54 amino acid residues 101 to 144) were randomized to NNN, where N could be A, G, C, or T, yielding a set of input libraries with 880 different single amino acid mutations, not including mutants in which the codons were changed to stop codons. All functional *rad54* variants were identified based upon their ability to support growth of a *rad54Δ* strain on media containing 0.015% MMS (Figure 3A). The resulting pool of survivors were analyzed by next generation DNA sequencing. Fold-enrichment for each variant was calculated by comparing the relative abundance of each allele recovered from cells grown in the absence of MMS versus cells grown in the presence of MMS (Figure 3B, 3C & Table S3). All of the wild-type alleles were enriched in the survivor pool, providing internal positive controls demonstrating that the deep mutagenesis screen could recover the *S. cerevisiae* wild-type *RAD54* alleles (Figure 3B). In contrast, alleles bearing stop codons were depleted, providing internal negative controls confirming that full-length Rad54 was necessary for cell growth on MMS (Figure 3B). Notably, with the exception of the FxxP motif, the functional landscape of the Rad51 interaction motif of Rad54 was largely tolerant of single point mutations, suggesting that no single amino acid residue within the motif was absolutely essential, although there were many instances where particular amino acid residues were not tolerated at some position (Figure 3B-D). For example, in agreement with the point mutation data, the L127A and L127D *rad54* mutants were 9-fold and 1,679-fold depleted in the deep mutagenic screen compared to wild-type *RAD54*. In addition, the deep mutagenic screen showed that although proline was highly preferred within the FxxP motif, mutations at residue P134 to alanine or glycine were also tolerated, which agreed with the point mutation data (Figure 3B-C & Figure S4). Lastly, comparison of the results of the deep mutagenesis to the sequence conservation of the Rad51 interaction motif based upon alignment of Rad54 sequences from 4,836 specifies shows good agreement between the two data sets and affirms that the phenylalanine and proline residues of the FxxP were the among the two most important residues with respect to Rad54 functional interactions with Rad51 (Figure 3D).

### Predicted structure of Hed1 bound to Rad51

Hed1 is a small (162 amino acid residues) meiosis-specific protein found in budding yeast that downregulates Rad51 activity in meiosis by blocking its interactions with Rad54 (Busygina et al., 2012; Busygina et al., 2008; Tsubouchi and Roeder, 2006). Based upon our previous single molecule assays, we have proposed that Hed1 acts as a competitive inhibitor of Rad54 binding and, given this model, we predicted that Hed1 would likely interact with the same binding surface on Rad51 as Rad54 (Crickard et al., 2018). As with Rad54, we used bioinformatics and AlphaFold3 predictions to identify potential regions of Hed1 responsible for its interactions with Rad51 (Figure 4A). No known homolog of Hed1 has yet been identified in higher eukaryotes, but Hed1 itself is conserved among the budding yeast and its CTD region contains the most highly conserved portion of the protein (Figure 4A-B). AlphaFold3 modeling of Hed1 alone predicts that the protein is largely unstructured (Figure S5A-S5B). However, modeling of Hed1 bound to Rad51 suggests that the CTD of Hed1 becomes highly structured upon binding to the Rad51-ssDNA nucleoprotein filament and its predicted binding site also overlapped with that of Rad54, (Figure 4C-D, Figure S5C). Notably, the predicted Rad51 interaction motif from Hed1 does not contain an FxxP motif and its sequence is entirely unrelated to the Rad51 interaction motif from Rad54. Therefore, based upon these findings, we considered the possibility that Hed1 amino acid residues 118 to 156 might represent a unique Rad51 interaction motif distinct from that found in Rad54.

**Figure 4.**
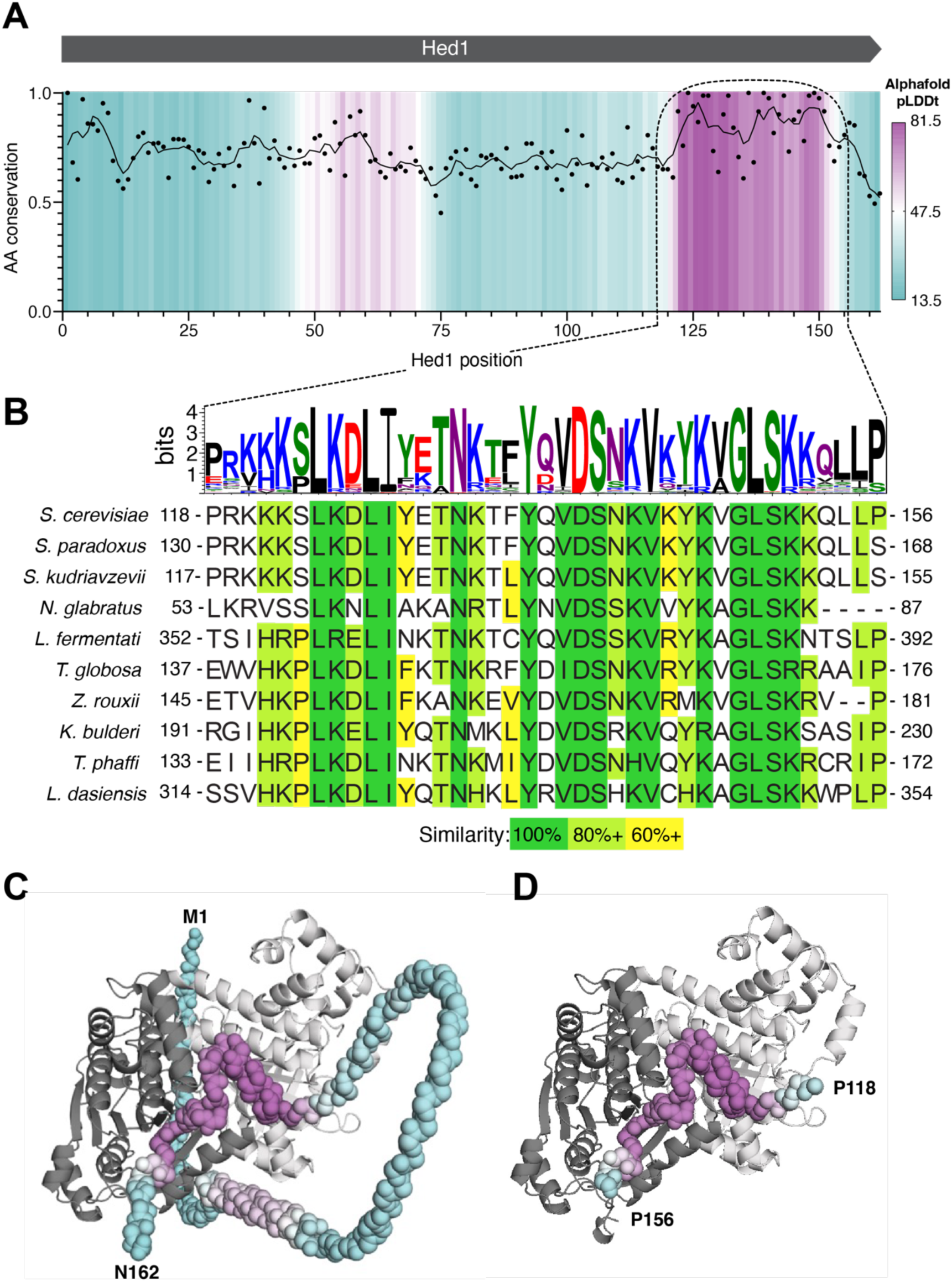
Predicted interaction interface between Rad51 and Hed1. **(A)** Plot of amino acid residue conservation and predicted AlphaFold3 structure (pLDDt; predicted local difference distance test) versus residue number for *S. cerevisiae* Hed1 is shown as in Figure 1A. **(B)** Sequence logo and sequence alignments for the region encompassing the highly conserved C-terminal region of Hed1 that is predicted to fold when bound to Rad51. **(C)** Predicted AlphaFold3 structure for full-length Hed1 (residues M1 to N162) and **(D)** Hed1 residues P118-P156 bound to two adjacent Rad51 monomers from within a Rad51 filament.

### Structure of the Hed1-bound Rad51 presynaptic complex

We obtained a 2.8 Å resolution CryoEM structure of the 39 amino acid residue Hed1 peptide (residues P118 to P156) bound to the Rad51-ssDNA filament and 33 of these residues were visible within the structure (residues K121 to L153; Figure 5A-5B, Figure S6-S7, & Table S1). Similar to Rad54, the Rad51 binding motif from Hed1 bridges two adjacent Rad51 monomers within the Rad51 filament corresponding to a buried surface area of 1,494.4 Å^2^ (Figure 5C). The ability of Hed1 to interact with two adjacent Rad51 monomers provides a structural explanation for the observation that Hed1 binding stabilizes Rad51 filaments (Busygina et al., 2012). The N-terminal most portion of the Hed1 peptide is comprised of a short loop made up of residues P118 to S123, followed by a three-turn alpha helix made up of residues L124 to F135, and it ends with a short loop region comprised of residues Y136 to P156 (Figure 5C-E). Most of the Hed1 amino acid residue contacts from this region involve a single alpha helix within the Rad51 protomer facing the 5’ end of the ssDNA (Figure 5D). These include Hed1 residues K122, T131, T134 and Y136 which are within hydrogen bonding distance of Rad51 residues E271, Q267, and H257, and Hed1 residue L124 which is involved in hydrophobic interaction with Rad51 residue A248(Figure 5D). Three additional Hed1 residues within this region contact the second Rad51 protomer: K125 makes contacts with Rad51 residue D149; residue I128 is involved in hydrophobic interactions with Rad51 residues F144 and V145; and residue N132 which makes hydrogen bond contacts with the backbone amino and carboxyl atoms of Rad51 residue F144 (Figure 5F). The C-terminal most portion of the Hed1 peptide is comprised of an extended loop-like region that interacts with the second Rad51 protomer via electrostatic and hydrophobic contacts (Figure 5E). The electrostatic contacts include Hed1 residues Q137, K146 and K152, which contact Rad51 residues R138, E108 and E156, respectively (Figure 5E). Hydrophobic contacts include Hed1 residues V138, V143, Y145 and L149, which interact with a hydrophobic pocket present on the surface of Rad51 comprised of residues A137, V140, P141, M142, G143, F144, G318, V319 and A320 (Figure 5E).

**Figure 5.**
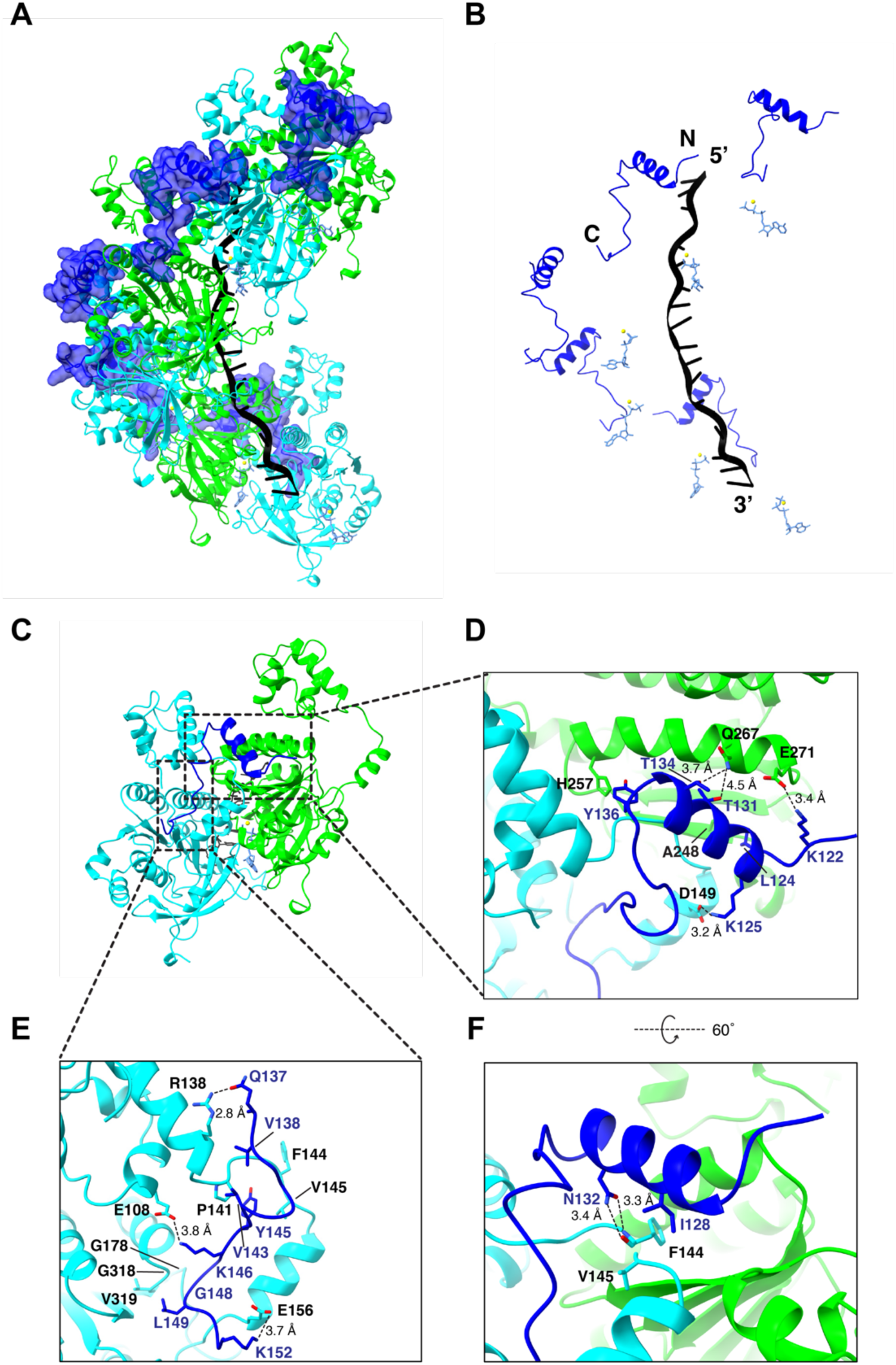
Structure of the Rad51 interaction motif of Hed1 bound to the Rad51-ssDNA filament. **(A)** CryoEM reconstruction of the Rad51-ssDNA filament bound by a peptide encompassing Hed1 residues P118 to P156. Rad51 subunits are shown in alternating colors (cyan and green) and the Hed1 peptide is shown in blue. **(B)** Structure in which the Rad51 proteins have been removed to highlight the relative locations of the Hed1 peptide, Rad51-bound ssDNA and ATP molecules. **(C)** Structure showing that the Hed1 peptide (blue) is bound to two adjacent Rad51 subunits (the 5’ facing subunit is shown in green and the 3’ facing subunit is shown in cyan). **(D-F)** Details of the molecular interactions between Hed1 (blue) and the two adjacent Rad51 subunits.

Interestingly, four Hed1 mutations have previously been identified that disrupt its interactions with Rad51, which include I128M, T131P, N132S, and N132I (Busygina et al., 2012). Our structure shows that I128 is involved in a hydrophobic interaction Rad51 residues F144 and V145, and the longer methionine side chain would likely induce a steric clash with these residues; and the Hed1 T131P mutation would abolish a hydrogen bonding interaction with Rad51 residue Q267 and disrupt helical secondary structure. The N132S and N132I mutations likely disrupt interactions with the backbone amino and carboxyl atoms of Rad51 residue F144. Another Hed1 mutant, Hed1 Δ114-122, can bind to Rad51 but its binding interaction is weakened sufficiently that it can no longer block the binding of Rad54 (Busygina et al., 2012; Crickard et al., 2018). Residues 114-117 lie just outside of the region that we define as the Rad51 interaction motif of Hed1, residues 118 to 120 are not visible in the structure, and residue 121 does not appear to interact with Rad51. However, residue K122 interacts with Rad51 residue E271, and disruption of this contact may contribute to the weakened binding observed for the Hed1 Δ114-122.

Most notably, the Hed1 peptide binds across the same surface of Rad51 as does Rad54, albeit through an entirely different mechanism, and it is in fact bound in the complete opposite orientation to that observed for Rad54 (c.f. Figure 2 & Figure 5). The Hed1 and Rad54 binding sites overlap with one another, such that the binding of Hed1 and Rad54 would be mutually exclusive, providing a clear structural explanation for prior single molecule studies showing that Hed1 acts as a competitive inhibitor of Rad54 binding (Crickard et al., 2018).

### Genetic analysis of the Rad51 interaction motif from Hed1

To further assess the biological relevance of the Hed1 contacts observed in the CryoEM structure, we adapted an assay in which the expression of Hed1 during mitosis causes cells to become sensitized to Rad51-mediated repair of MMS-induced DNA damage (Tsubouchi and Roeder, 2006). As expected, expression of wild-type Hed1 in mitotic cells resulted in sensitivity to MMS, whereas the empty vector control did not (Figure S8A). We also tested the previously published *hed1* mutants I128M, T131P, and N132S, all of which allowed for cell growth on MMS (Figure S8A), consistent with previously published results (Busygina et al., 2012). In striking contrast to Rad54, single alanine point mutations in most residues that make specific contacts between Hed1 and Rad51 resulted in the loss of Hed1-mediated inhibition of cell growth, including residues K122, L124 K125, T131, Y136, V138, V143, K146, L149, and K152 (Figure S8A). The exceptions were T134A, which exhibited a phenotype comparable to wild-type Hed1, and Q137A, which yielded a mild phenotype consistent with partial inactivation of Hed1 activity (Figure S8A). Notably, all *hed1* variants tested that contained multiple mutations exhibited null phenotypes (Figure S8B). Together, these genetic assays provide support for the biological relevance of the protein-protein interface observed on our CryoEM structure of the Rad51-Hed1 complex.

## DISCUSSION

The Rad51/RecA family of proteins is responsible for the DNA pairing reactions that take place during homologous recombination and given their central role in recombination these proteins are subject to both positive and negative regulation by a number of different auxiliary factors. Rad54 is a key example of a positive regulatory factor that strongly stimulates the strand exchange activity of Rad51, and in budding yeast the importance of Rad54 is revealed by the fact that deletion of the *RAD54* gene is just as deleterious to cell survival as is the loss of *RAD51* when cells are subjected to DNA damage. Here we have identified the Rad51 interaction motif within Rad54, and we have described the structural basis for Rad54 interactions with the Rad51 presynaptic complex. We have also provided the structural basis for how the interaction between Rad51 and Rad54 is negatively regulated by the meiosis-specific protein Hed1 allowing for meiotic recombination to be driven by meiosis-specific recombinase Dmc1. Together, our findings provide atomic-level insights into the spatial and temporal interplay between Rad54, Hed1, Rad51 and Dmc1 that takes place during the transition from mitotic to meiotic recombination in *S. cerevisiae*.

### Rad54 interactions with the Rad51 presynaptic complex

The Rad54 binding sites on Rad51 span the entire length of the presynaptic complexes suggesting that Rad54 could, in principle, act in approximately 1:1 stoichiometry with Rad51 (Figure 2). This finding is consistent with previous *in vitro* biochemical studies which suggested that an optimal ratio of one molecule of Rad54 for each molecule of Rad51 during strand invasion (Mazin et al., 2000). However, recent studies using GFP-tagged proteins in live *S. cerevisiae* cells suggest that Rad54 functions at sub-stoichiometric concentrations compared to Rad51 (A. Taddei, personal communication), which agrees with prior single molecule studies showing that the homology search can be supported by sub-stoichiometric levels of Rad54 (Crickard et al., 2020b). These considerations raise the interesting question of whether Rad51 in cells might be bound by Rad54 monomers at locations distributed randomly along the length of the presynaptic filament, or whether Rad54 might form localized clusters of more well-defined higher order complexes.

Notably, the Rad54 binding pocket spans two adjacent Rad51 monomers within the Rad51 presynaptic complex (Figure 2C), which may help explain two important influences of Rad54 on Rad51 behavior. First, the ability of Rad54 to bridge two adjacent Rad51 monomers within the filament provides a plausible explanation for how Rad54 stabilizes the presynaptic complex (Mazin et al., 2003b; Meir et al., 2022). Thus, Rad54 may act as a molecular “staple” that helps maintain nucleoprotein stability. Second, the existence of a composite binding pocket comprised of two adjacent Rad51 monomers provides a straightforward structural explanation for why Rad54 is recruited to sites of DNA damage only after the initial recruitment of Rad51 (Lisby et al., 2004), in that the composite Rad54 binding pocket would only exist after the assembly of Rad51 filaments onto the ssDNA overhangs present at processed double strand breaks. Thus, the existence of a composite binding surface for Rad54 on the surface of the Rad51-ssDNA presynaptic complex likely serves as an important regulatory mechanism for controlling the timing of Rad54 recruitment to the sites of DNA damage.

### Conservation of the Rad51-Rad54 binding mechanism

Our CryoEM data shows that contacts with Rad51 span a region within *S. cerevisiae* Rad54 encompassed by amino acid residues T101 to L144. Interestingly, with the exception of the phenylalanine and proline residues within the FxxP motif, our deep mutagenic analysis reveals that the functional landscape of the Rad51 interaction motif of Rad54 is highly tolerant to single amino acid variants, which is also in good agreement with the sequence conservation data (Figure 3). Importantly, the essential phenylalanine and proline of the Rad54 FxxP sequence motif is conserved from yeast to humans as is the corresponding binding pocket on the Rad51 filament (Figure S9), suggesting that the mode of interaction defined by our CryoEM structure of the Rad54 peptide bound to the Rad51 presynaptic complex may be broadly conserved. Consistent with this conclusion, AlphaFold3 predictions for human RAD54 provide further evidence for a conserved mode of interaction, with a similar predicted backbone fold and hydrophobic contacts (Figure S9A-S9C). This hypothesis is further supported by docking of the Rad54 peptide into our previous structure of *S. cerevisiae* Dmc1 (PDB: 9D4N; (Petassi et al., 2024)), which reveals that all of the major contacts are maintained (Figure 6B). In addition, the FxxP binding pocket is also conserved in both yeast and human Dmc1 (Figure S9C-S9D), strongly suggesting that the same mode of interaction may be utilized for Rad54 interactions with Dmc1 (*see below*).

**Figure 6.**
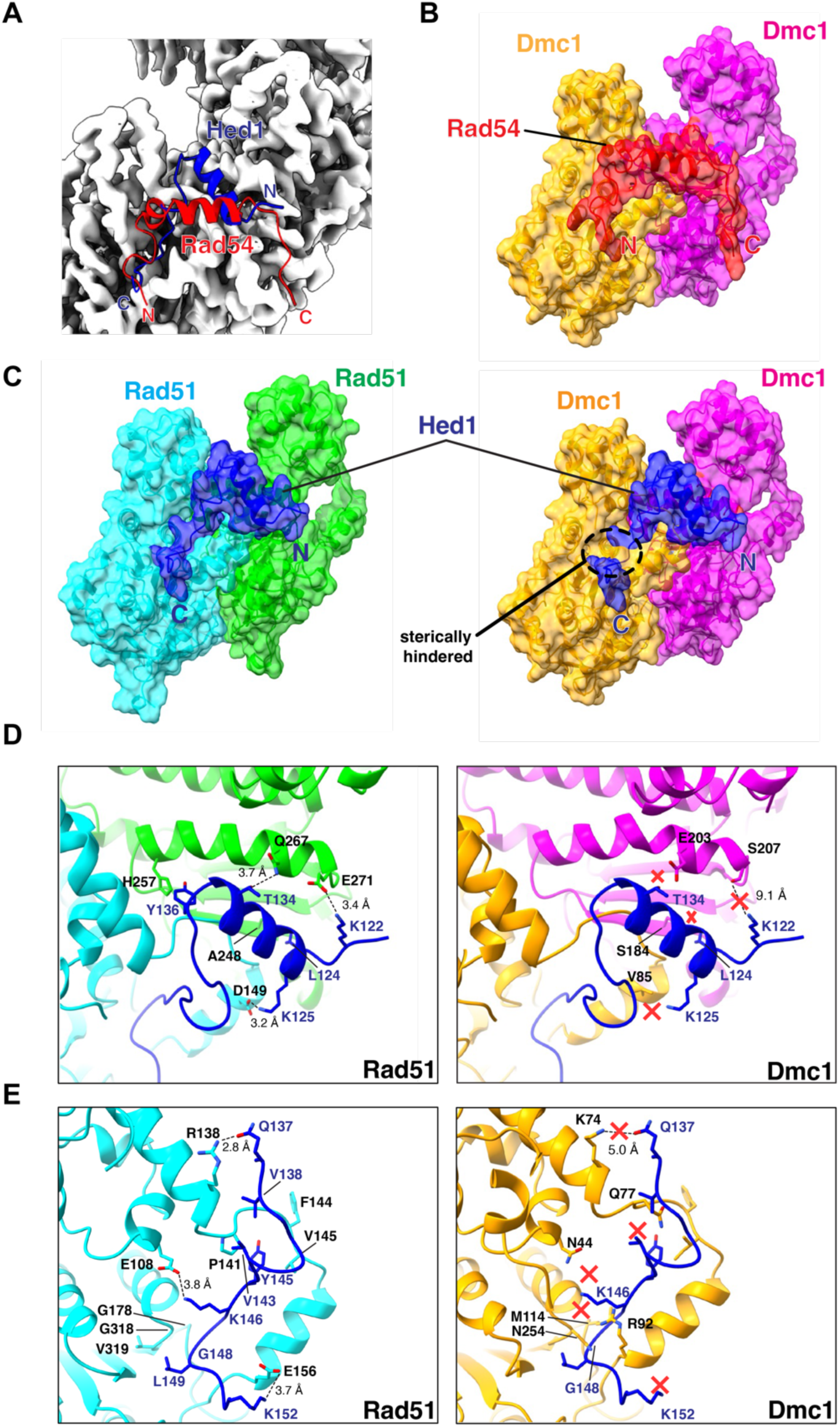
Structural basis for the mechanisms of Rad51 regulation in meiosis. **(A)** Model showing that the Rad51 interaction motifs from Rad54 (red) and Hed1 (blue) overlap with one another when bound to Rad51 (gray) but are oriented in the opposite directions relative to their N- and C-termini. **(B)** Illustration showing that the Rad51 interaction motif peptide from Rad54 (red) can be docked onto the structure of *S. cerevisiae* Dmc1 (adjacent subunits shown in orange and magenta). **(C)** Side-by-side comparison of the Rad51 interaction motif from Hed1 (blue) bound to either Rad51 (adjacent subunits shown in cyan and green) or docked onto Dmc1 (adjacent subunits shown in orange and magenta). **(D-E)** Side-by-side comparisons of the molecular contacts between Hed1 and Rad51 or Hed1 and Dmc1. Hed1 contacts that are predicted to be disrupted with Dmc1 are highlighted with a red “x”.

Interestingly, human RAD51 binds to the tumor suppressor protein BRCA2 via the RAD51-binding TR2 motif located within the C-terminus of BRCA2 (Appleby et al., 2023; Davies and Pellegrini, 2007; Dunce and Davies, 2024; Esashi et al., 2007; Kwon et al., 2023; Sharan et al., 1997). This interaction also involves an FxxP motif found within BRCA2 (Appleby et al., 2023; Dunce and Davies, 2024) and the binding site for the TR2 motif overlaps with the same site where we predict that human RAD54 will interact with RAD51 (Figure S9E). Thus, accumulating evidence suggests that multiple factors may interact with same binding cleft on the surface of the Rad51-ssDNA presynaptic complex, which implies the possible requirement for different regulatory mechanisms to control the spatial distribution and temporal association of required factors at various stages of the homologous recombination, such as Rad51 filament assembly, the homology search, strand invasion, and Rad51 filament disassembly.

Rdh54 is a closely related homolog of Rad54, and it also participates in homologous recombination (Dresser et al., 1997; Klein, 1997; Shinohara et al., 1997). Known attributes of Rdh54 include stimulating D-loop formation *in vitro* (Nimonkar et al., 2012; Petukhova et al., 2000), the ability to remove Rad51 and Dmc1 from dsDNA (Chi et al., 2006; Holzen et al., 2006; Shah et al., 2010b), and acting together with Rad54 to enhance the stability of Rad51 presynaptic filaments (Meir et al., 2022), promoting template switching during break-induced replication (BIR)(Tsaponina and Haber, 2014), playing a greater role in interhomolog recombination compared to Rad54 (Chi et al., 2006; Klein, 1997; Shinohara et al., 1997). Notably, Rdh54 and Rad54 both bind to the Rad51 presynaptic complex through their disordered N-terminal domains, but they interact through different binding mechanisms that are not mutually exclusive (Crickard et al., 2020a). The evidence supporting this idea is that (*i*) Rad54 and Rdh54 can both simultaneously interact with the presynaptic complex without interfering with one another, (*ii*) Hed1 selectively inhibits Rad54 binding to Rad51 but has no effect on Rdh54 binding, but (*iii*) Hed1 can be made to selectively inhibit Rdh54 just by swapping the NTDs between Rad54 and Rdh54 (Crickard et al., 2020a). Therefore, we predict that Rdh54 may display a new mode of interaction with the Rad51-ssDNA presynaptic complex that will be distinct from that of both Rad54 and Hed1.

### Structural basis for the regulatory control of Rad51 and Dmc1 in meiosis

Rad51 is the only recombinase that is expressed in mitosis and during HR-dependent DNA repair it drives strand invasion primarily between sister chromatids. Entry into meiosis coincides with the expression of the meiosis-specific recombinase Dmc1 (Bishop et al., 1992) and the meiosis-specific protein Hed1, which downregulates Rad51 strand exchange activity (Tsubouchi and Roeder, 2006). As a consequence, Dmc1 is considered to be the active recombinase responsible for strand exchange between homologous chromosomes in meiosis (Brown and Bishop, 2014; Cloud et al., 2012). Our data provide a structural explanation for how this regulatory transition takes place in budding yeast. Our CryoEM structures show that Hed1 and Rad54 bind to the same surface cleft on Rad51 such that their binding interactions would be mutually exclusive (Figure 6A), and we have previously shown that these binding interactions are so tight that they show no evidence for reversible dissociation *in vitro* on time scales that might be considered biologically relevant (Crickard et al., 2018).

In striking contrast to Rad54, which interacts with both Rad51 and Dmc1, Hed1 is highly specific for just Rad51 and does not bind to Dmc1 (Crickard et al., 2018). This binding specificity ensures that Hed1 can selectively downregulate Rad51 strand exchange activity in meiosis without affecting the strand exchange activity of Dmc1. Our structural studies provide a plausible explanation for this specificity by showing that there would be extensive steric clashes between the C-terminal region of the Hed1 peptide and Dmc1 if the same binding mode was preserved (Figure 6C). There are also numerous other Rad51 residues that make contacts with Hed1 that are different in Dmc1 such that they disrupt essential protein-protein contacts (Figure 6D & 6E). For example, Rad51 residues E108, E156, D149 and Q267 are instead N44, R92, V85, and E203 in Dmc1, all of which result in the disruption of specific residue interactions (Figure 6D & 6E). Rad51 residue R138 is K74 in Dmc1 and would be positioned 2 Å further away from Hed1 residue Q137, perhaps weaking this interaction. Rad51 residue P141 aligns with residue Q77 in Dmc1, resulting in the loss of a hydrophobic interaction (Figure 6E). Rad51 residue E271, which interacts with Hed1 residue K122, is equivalent to S207 in Dmc1 and is positioned too far away to form a hydrogen bonding interaction (Figure 6D). Lastly, Rad51 residues G178 and G318 are instead M114 and N254 in Dmc1, which would sterically clash with Hed1 (Figure 6E).

Notably, seven of the Hed1 contacts with Rad51 involve amino acid residues that are specifically conserved only within the Rad51 lineage of the Rad51/RecA family of proteins and are different residues within the Dmc1 lineage (*manuscript in preparation*). These include *S. cerevisiae* Rad51 residues E108, P141, E156, G178, A248, M268 and G318, which correspond to *S. cerevisiae* Dmc1 residues N44, Q77, R92, M114, S184, E204, and N254, respectively. Interestingly, chimeric *rad51* mutants harboring these Dmc1 lineage-specific amino acids (E108N, P141Q, E156R, G178M, A248S, M268E and G318N) all yield biologically nonfunctional proteins incapable of supporting cell growth in the presence of MMS (*manuscript in preparation*). Moreover, the fact that these residues are broadly conserved within the Rad51 lineage, whereas Hed1 thus far has only been identified in yeast species, suggests the possibility that Hed1-like meiosis-specific regulatory proteins, which bind to Rad51 but not Dmc1, may also exist in other eukaryotes, thus allowing Dmc1 to act as the dominate recombinase in meiosis (Davies and Pellegrini, 2007; Esashi et al., 2007; Kwon et al., 2023; Sharan et al., 1997).

### Regulation of Rad54 binding via post-translational modification

In *S. cerevisiae*, an additional level of regulatory control is exerted over Rad54 during meiosis via Mek1-dependent phosphorylation of T132 (Niu et al., 2009), which may weaken interactions with Rad51 by adding a phosphate group immediately adjacent to the Rad54 FxxP motif that is in close proximity to the protruding acid patch (PAP) of Rad51 (Figure 2F). Although Rad54-Rad51 interaction mechanism may be broadly conserved (Figure S9), its regulation through Mek1-dependent phosphorylation of residue T132 may be specific to budding yeast given that threonine 132 is not conserved and is instead replaced with lysine in many higher eukaryotes (Figure 1C).

## Conclusion

We present two new structures of the Rad51 presynaptic filament bound to peptide fragments from Rad54 and Hed1, both of which play important regulatory roles in homologous recombination. We show that these regulatory factors interact with the same surface on the Rad51 presynaptic complex but utilize distinct binding mechanisms to allow for differential regulation of Rad51 strand exchange activity during mitosis and meiosis. This particular binding surface on Rad51 has now been implicated in numerous protein-protein interactions and competition between different accessory factors for this same site may reflect a broad theme in the regulatory control homologous recombination.

### Limitations of the study

Although we have provided insights into the interplay between Rad51, Dmc1, Rad54 and Hed1, several important questions remain to be answered. In particular, our structural studies were conducted with peptides encompassing the Rad51 interaction motifs of Rad54 and Hed1 rather than full-length Rad54 or full-length Hed1, so we may be missing evidence of more extensive interactions involving residues flanking the Rad51 interaction motifs. Similarly, we do not yet know how full-length Rad54 or full-length Hed1 are organized with respect to the Rad51-ssDNA filaments. Lastly, we rely upon AlphaFold3 predictions for our discussion of *S. cerevisiae* Dmc1, human RAD51, and human DMC1, so the models we present with respect to these proteins are currently predictive but should not be taken as direct experimental evidence.

## RESOURCE AVAILABILITY

### Lead contact

Requests for further information and resources should be directed to and will be fulfilled by the lead contact, Eric C. Greene.

### Materials availability

Plasmids and strains generated in this study are available upon request, which should be directed to the lead contact, Eric C. Greene.

### Data and code availability

- The CryoEM structures of Rad54-Rad51-ssDNA and Hed1-Rad51-ssDNA have been deposited in the Research Collaboratory for Structural Bioinformatics (RSCB) Protein Data Bank (PDB) with accession codes: 9E6L and 9E6N.
- The CryoEM density maps of Rad54-Rad51-ssDNA and Hed1-Rad51-ssDNA have also been deposited in the Electron Microscopy Data Bank (EMDB) with accession codes: EMD-47572 and EMD-47573.
- All original code has been deposited in Github and is publicly available at the following URL: https://github.com/michaeltpetassi/Rad54Hed1_2025
- Deep sequencing data have been deposited at the NCBI Sequence Read Archive (SRA) and are publicly available via the BioProject ID: PRJNA1235687.
- Any additional information required to reanalyze the data reported in this paper is available from the lead contact upon request.

## Supporting information

Spreadsheets of data related to the manuscript

## ACKNOWLEDGEMENTS

We thank Andreas Hochwagen, Rodney Rothstein, Lorraine Symington and Luke Berchowitz for yeast strains and access to equipment. We thank members of the Greene laboratory for critically reading the manuscript. This research was funded by NIH grant R35GM118026, R01CA236606 and P01CA275717, and an NSF Grant MCB-1817315 (to E.C.G.). S.K. and K.M. were partially supported by the Columbia University Summer Undergraduate Research Fellowships (SURF) program and the Columbia College Summer Funding Program. This research was funded in part through the NIH/NCI Cancer Center Support Grant P30CA013696 and used the Genomics and High Throughput Screening Shared Resource. The CryoEM was in part supported by the National Cancer Institute’s National Cryo-EM Facility at the Frederick National Laboratory for Cancer Research under contract 75N91019D00024, some of the work was performed at the National Center for CryoEM Access and Training (NCCAT) and the Simons Electron Microscopy Center located at the New York Structural Biology Center, supported by the NIH Common Fund Transformative High Resolution Cryo-Electron Microscopy program (U24 GM129539,) and by grants from the Simons Foundation (SF349247) and NY State Assembly, and some of this work was performed at the Columbia University Cryo-Electron Microscopy Center.

## AUTHOR CONTRIBUTIONS

Y.S. purified proteins and solved all CryoEM structures. M.T.P. performed the bioinformatic analysis, deep mutagenic screens and genetic analysis of Rad54 and Hed1 mutants. Y.S. and M.T.P both conducted the AlphaFold3 analyses. S.Y.K. assisted with Rad51 protein purification. A.J. and R.M. conducted genetic analysis of Rad51 PAP mutants. K.M. assisted with the genetic analysis of Rad54 and Hed1 mutants. V.B.R. assisted with the design of all genetic experiments. U.R. conducted an initial genetic analysis of the Rad51 PAP mutants and performed the PAP mutant D-loop assays. All authors assisted with preparation of the manuscript.

## DECLARATIONS OF INTERESTS

The authors declare no competing interests.

## STAR METHODS

### Bioinformatics

For Rad54, *S. cerevisiae* S288C RAD54 (NP_011352) and *S. cerevisiae* S288C Rdh54 (NP_011106) were used as queries for blastp searches with an E-value threshold of 1e-60 against the NCBI nr protein database accessed in July 2023 (Sayers et al., 2022). Sequences shorter than 650 amino acids or longer than 1300 amino acids were discarded, resulting in 10,401 unique sequences. Sequences were clustered to 60% identity using cd-hit (Fu et al., 2012) resulting in 993 clusters, representatives aligned with MUSCLE (Edgar, 2004), and a similarity tree was built using Genieous Prime 2023.1 https://www.geneious.com). Sequences clustered into three clades, corresponding to Rad54, Rdh54, and a small group of distantly related helicases XRCC6. The Rad54 clade (4,855 sequences) was extracted, clustered to 80% identity using cd-hit (898 clusters) and aligned with MUSCLE. For Hed1, *S. cerevisiae* S288C HED1 (NP_001035220) was used as query for blastp search with an E-value threshold of 0.05 against the NCBI nr protein database accessed in June 2024 (1). The 87 resulting sequences were aligned with MUSCLE.

### Protein purification

*S. cerevisiae* Rad51 was overexpressed in *E. coli* BL21 (DE3) Rosetta2 cells transformed with a plasmid encoding 6XHis-SUMO-ScRad51. Cells were grown in 2L LB media containing 100 μg/ml carbenicillin and 35 μg/ml chloramphenicol at 37°C until OD600 = 0.6, induced with 0.5 mM IPTG and grown for 3 hours at 37°C. The cell paste was suspended in 50 ml lysis buffer (50 mM Tris-HCl [pH 7.5], 10% glycerol, 1M NaCl, 1mM DTT, 0.5 mM PMSF, 15 mM imidazole, 0.1% tween 80, 1 protease inhibitor tablet) and lysed by sonication. The lysate was centrifuged at 35,000 rpm for 45 mins followed by precipitating the supernatant with 12g ammonium sulfate for 1 hour. The precipitate was spun down at 10,000 rpm for 30 mins. The pellet was dissolved in 50 ml binding buffer (25 mM Tris-HCl [pH 7.5], 10% glycerol, 200 mM NaCl, 0.1% Triton X-100, 15 mM imidazole, 5 mM beta-mercaptoethanol) and applied to the 5 ml HisPur^TM^ Ni-NTA resin (Thermo Fisher Scientific) equilibrated with the same binding buffer. The protein was eluted with 10 ml elution buffer (25 mM Tris-HCl [pH 7.5], 10% glycerol, 200 mM NaCl, 200 mM imidazole, 0.1% Triton X-100). SUMO protease was added to the elution, followed by dialyzing for 16 hours at 4 °C in dialysis buffer (50 mM Tris-HCl [pH 7.5], 200 mM NaCl, 10% glycerol, 15 mM imidazole, 1 mM DTT). The sample was re-applied to the 5 ml Ni-NTA resin equilibrated with the binding buffer and flow-through was collected and concentrated to 30 μM. The protein was flash-frozen in liquid nitrogen and stored at –80 °C.

The Rad51 triple mutant protein (D239A, D241A, D242A) was purified in the same way as wild-type Rad51, except it was dialyzed and stored in buffer with 500 mM NaCl to prevent aggregation. In brief, 6xHis–SUMO–Rad51-D239A, D241A, D242A was overexpressed in *E. coli* BL21 (DE3) Rosetta2 cells at 37 °C to an OD_600_ of 0.4–0.6. Expression was induced by addition of 0.5 mM IPTG for 3 h at 37°C. Cells were harvested and stored at –80 °C. Cells were lysed in Cell Lysis Buffer (30 mM Tris–HCl [pH 8.0], 1 M NaCl, 10% glycerol, 10 mM imidazole, 1 mM DTT, 200 mM PMSF, 10% NP-40 and protease inhibitor cocktail (Roche, Cat. No. 05892953001)) by sonication for 10 (s) on and 30 (s) off for a total time of 2 (min). The lysate was clarified by ultracentrifugation at 100,000 x g for 45 min at 4 °C and proteins precipitated with 40% ammonium sulfate. The precipitated proteins were redissolved in Cell Lysis Buffer and bound to 1.5 mL of pre– equilibrated Ni–NTA resin. The resin was then washed 3X with CLB and eluted in CLB + 200 mM imidazole. The eluted fraction was mixed with 400 units of the SUMO protease Ulp1 (Sigma–Aldrich, Cat. No. SAE0067–2500UN) and dialyzed overnight at 4 °C into Rad51 buffer (30 mM Tris–HCl [pH 8.0], 500 mM NaCl, 1 mM EDTA, 10% Glycerol, 10 mM imidazole). The 6xHis–SUMO tag and SUMO protease were removed by passing the dialyzed proteins over a second 1.5 mL Ni–NTA column. The purified Rad51-D239A, D241A, D242A protein was stored at –80 °C in single use aliquots.

For use in D-loop assays, full-length Rad54 was purified as previously described (Crickard et al., 2020b). A protease deficient yeast strain was transformed with GST– tagged Rad54 on a 2–micron plasmid under the control of the Gal1 promoter. Cells were grown in Yeast Nitrogen base (–URA) plus 3% Glycerol and 2% lactic acid. When the cells reached an OD_600_ of 1.5, expression was induced by the addition of 2% galactose for 6 hours. Cells were harvested and stored at –80 °C. Cell pellets were re-suspended in Rad54 re-suspension buffer (30 mM Tris–HCl [pH 7.5], 1 M NaCl, 1 mM EDTA, 10% glycerol, 10 mM BME (β-mercaptoethanol), Protease inhibitor cocktail (Roche Cat. No. 05892953001) and 2 mM PMSF. Cells were disrupted by manual bead beating, and the lysate was clarified by ultracentrifugation at 100,000xg for 1 hour. The lysate was fractionated by ammonium sulfate (AS) precipitation. AS was gradually added with mixing to a final concentration of 20% followed by centrifugation at 10,000Xg for 10 minutes. The supernatant was discarded, and the AS concentration was raised to 50% followed by centrifugation at 10,000x g for 10 min. The protein pellet was re–suspended in PBS (phosphate buffered saline) plus 1 M NaCl and 10 mM BME. The resulting re– suspended protein was then bound to pre–equilibrated GST resin in batch for 1 hour at 4 °C. The GST resin was washed with PBS plus 1 M NaCl and washed again with PBS plus 500 mM NaCl. The protein was then eluted with 20 mM glutathione in PBS plus 500 mM NaCl. The peak fractions were pooled and the applied to a Sephacryl S–300 High Resolution gel filtration column (GE Healthcare, Cat. No. 17–0599–10) pre–equilibrated with Rad54 SEC buffer (30 mM Tris-HCl [pH 7.5], 500 mM NaCl, 1 mM EDTA, 10 % glycerol, and 10 mM BME. The protein eluted in two peaks, one peak occurred outside of the exclusion volume of the column and was discarded. The second peak eluted near the expected MW of a Rad54 monomer and was collected. The monomeric Rad54 peak was pooled and dialyzed against Rad54 SEC buffer plus 50 % glycerol and stored in at –80°C in single use aliquots.

### AlphaFold3 structure predictions

Alphafold3 predictions were performed using the Google DeepMind Alphafold3 Server (https://golgi.sandbox.google.com/)(Abramson et al., 2024) with the following molecules - recombinase (*S. cerevisiae* Rad51 UniProt P25454, *H. sapiens* Rad51 UniProt Q06609, *S. cerevisiae* Dmc1 UniProt P25453, or *H. sapiens* Dmc1 UniProt Q14565), poly(dT) ssDNA, ATP, Mg2+, and one interacting protein (*S. cerevisiae* Rad54 UniProt P32863, *S. cerevisiae* Hed1 UniProt Q03937, *H. sapiens* RAD54L UniProt Q92698).

### CryoEM sample preparation

Peptides sequences were as follows: Rad54 peptide, 44 amino acids, derived from residues 101-144 of *S. cerevisiae* Rad54: 101-T K R R K D A L S A Q R L A K D P T R L S H I Q Y T L R R S F T V P I K G Y V Q R H S L-144; and the Hed1 peptide, 39 amino acids, derived from residues 118-156 of *S. cerevisiae* Hed1: 118-P R K K K S L K D L I Y E T N K T F Y Q V D S N K V K Y K V G L S K K Q L L P-156. Peptides were synthesized by Genscript (>95% purity) and were resuspended in phosphate buffered saline (PBS) + 1 mM DTT to a final concentration of 400 µM for the Rad54 peptide or 200 µM for the Hed1 peptide.

To assemble the nucleoprotein filaments, Rad51 (5 μM) and a 96-mer ssDNA (0.25 μM) were preincubated for 10 min at 30 °C in HR buffer (30 mM Tris [pH 7.5], 20 mM MgCl_2_, 50 mM KCl, 1 mM DTT, 2 mM ATP). To form the Rad51-Rad54 peptide and Rad51-Hed1 peptide complexes, an 8-fold molar excess (40 μM) of the either the Rad54 peptide or the Hed1-peptide was added to the pre-assembled Rad51-ssDNA nucleoprotein filaments and reactions were further incubated for an additional 10 min at 30 °C. A 3.5 μL aliquot of these samples was applied to glow-discharged UltrAuFoil R 0.6/1 Au300 grid for the Rad54 peptide and AuFlat 1.2/1.3 grid for Hed1 peptide, blotted for 5 seconds with a force 3, and plunge-frozen in liquid ethane using a Vitrobot Mark IV (FEI, USA) with 100% humidity at 4 °C.

### Cryo-EM data acquisition

Samples were initially screened using a Glacios (Thermo Fisher, 200 keV) at Columbia University Irving Medical Center. Grids selected for high-resolution data collection were imaged using a Titan Krios microscope (Thermo Fisher) operated at 300 keV and equipped with a K3 direct electron detector at the New York Structural Biology Center. For the Rad54 complex and the Hed1 complex, the magnification was 105K (pixel size = 0.844 Å/pixel) and 85K (pixel size = 1.083 Å/pixel) collected in electron counting mode, with a total dose of 59.51 e^-^/Å^2^ and 51.21 e^-^/Å^2^, respectively. Defocus range was −0.8 to −2.5 for both data sets.

### CryoEM data processing

Data was processed using CryoSPARC v4.3.1 (Punjani et al., 2017). For the Rad54 peptide complex, a total of 8,367 movies were collected and aligned using patch motion correction and patch CTF estimation (Rohou and Grigorieff, 2015). After manual inspection of micrographs for quality such as ice contamination, 246 micrographs were discarded, and 8,121 micrographs were used for downstream processing. Initially, 995,541 particles were picked and extracted with a box size of 256x256 followed by iterative 2D classification to remove junk/denatured and non-helical particles. The selected 434,806 particles were used for generating 2 *Ab initio* 3D density maps to generate an initial filament volume and clean out residual junk particles. 363,379 particles were used for the reconstruction and refinement of the final 3D map. The nominal resolution of the density map of 3.26 Å was estimated by 0.143 gold standard Fourier Shell Correlation (FSC) cut off (Figure S2).

For the Hed1 peptide complex, a total of 6,212 micrographs were collected and aligned using patch motion correction and patch CTF estimation. After manual inspection of micrographs for quality, 758 micrographs were discarded, and 5,454 micrograph was used for downstream processing. Initially, 1,184,183 particles were picked and extracted with a box size 256x256 followed by iterative 2D classification to remove junk/denatured and non-helical particles. The selected 895,334 particles were used for generating 2 *Ab initio* 3D density maps to generate an initial filament volume and clean out residual junk particles. 876,496 particles were used for the reconstruction and refinement of the final 3D map. The nominal resolution of the density map of 2.81 Å was estimated by 0.143 gold standard Fourier Shell Correlation (FSC) cut off (Figure S6).

### Structure refinement and analysis

Our previous structure of the Rad51-ssDNA filament (PDB: 9D46; (Petassi et al., 2024)) was used as a guide in building the atomic models. The Alphafold3 predicted peptide structures were used as a reference in manual fitting of the structures into the cryo-EM density map using Chimera(Pettersen et al., 2004). After initial rigid-body refinement in Phenix (Afonine et al., 2010), amino acid residue sidechains were manually inspected and corrected for fitting into the density map using Coot (Emsley and Cowtan, 2004). Fitted models were real space refined with the secondary structure and Ramachandran restraints in Phenix (Afonine et al., 2010).

Surface Area was calculated using PDBe PISA v1.52 ( http://www.ebi.ac.uk/pdbe/prot_int/pistart.html)(Krissinel and Henrick, 2007). The Rad51 dimer was considered as one structural unit and either the Rad54 or Hed1 peptide was considered as another structural unit for the calculation. All potential electrostatic interactions shown in the figures were verified using the FindHBond tool in UCSF Chimera with the geometric distance and angle constraints set to <4 Å and 110-180 degrees (Pettersen et al., 2004).

### Yeast transformations

Banked frozen yeast strains were streaked onto solid YPD media (1% yeast extract (Sigma-Aldrich, Cat No. 92144), 2% bacto-peptone (Sigma-Aldrich, Cat. No. 91249), 2% glucose (Sigma-Aldrich, Cat. No. G8270), 2% agar (Sigma-Aldrich, Cat. No. G8270) and grown for 2-3 days at 30 °C (all strain information is presented in Table S1). Cultures inoculated from single colonies were grown overnight in liquid YPD media and diluted 1:50 into fresh YPD to approximate OD_600_ = 0.4. After 4-hour incubation at 30 °C, cells were harvested by centrifugation at 3,000 rcf (Relative Centrifugal Force) for one minute, washed three times in at least 1/40 culture volume 0.1 M lithium acetate (Sigma-Aldrich, Cat. No. 517992) and resuspended in a final 1/125 culture volume 0.1 M lithium acetate. 50 µL competent cells were added to 360 µL transformation mix containing 33.33% PEG-3350 (Sigma-Aldrich, Cat. No. 202444), 0.1 M lithium acetate, 10 µg sheared salmon sperm DNA (Thermo Scientific, Cat. No. AM9680, incubated at 95 °C for 5 minutes then kept on ice until use), and up to 2 µg transforming plasmid. Mixes were incubated at 30 °C for 30 minutes with shaking, then incubated at 42 °C for 20 minutes without shaking. Cells were harvested at 3,000 rcf for 90 seconds, supernatant removed and resuspended in 100 µL sterile water before plating on synthetic dropout (SD) media (0.171% YNB (Sunrise Science Products, Cat. No. 1500), CSM lacking tryptophan (-Trp), leucine (-Leu) or uracil (-Ura) (Sunrise Science Products, Cat. No. 1007/1005/1004, recommended amount), 0.5% ammonium sulfate (Sigma-Aldrich, Cat. No. 517992), 2% glucose, 2% agar). Reactions were scaled up by 20-times in 50 mL conical tubes, as needed.

### Rad54 deep mutagenesis screen

*RAD54* and 500-bp upstream sequence was cloned into pRS415 (centromeric origin, *LEU2* marker) by generating three fragments by PCR amplification (PrimeSTAR Max DNA Polymerase, Takara Bio, Cat. No. R045B): two from *RAD54 S. cerevisiae* W303 genomic DNA covering +500-*RAD54* with an overlap region producing a silent mutation at K152 (A456G) to disrupt a 10-adenine tract with oligonucleotide primers MTP85/MTP135 and MTP86/MTP134; and a third PCR fragment from purified pRS415 with oligonucleotide primers VBR15/VBR16 (all oligonucleotide sequences are presented in Table S2). PCR products were digested with DpnI (20 units; NEB, Cat. No. R0176), assembled using In-Fusion Snap Assembly (Takara Bio, Cat. No. 638947) and transformed into Stellar competent *E. coli* cells (Takara Bio, Cat. No. 636763).

Mutant libraries of *pRS415*-*rad54* for deep mutagenesis screens were generated by single-fragment amplification of *pRS415-RAD54* with forward primer containing 9-bp mixed-base sequence at each 3-codon window and reverse primer generating 15-bp proximal sequence overlap for assembly protocol as above. A minimum of one million transformants were collected in LB supplemented with 100 μg/mL carbenicillin, diluted 1:20 in the same media (to OD_600_ of 2-3), and incubated at 37 °C with shaking for four hours before plasmids were extracted.

The *pRS415-rad54* mutant libraries were transformed into *S. cerevisiae* strain yECG46 as described above. After a 72-hour incubation at 30 °C, ≥1,000,000 transformants were collected, resuspended to a calculated OD = 100, and 100 µL were plated on SD –Leu supplemented with 0.015% MMS. After a 72-hour incubation at 30 °C, ≥10,000 colonies were collected. Plasmids were extracted from cell populations before and after MMS selection using Zymoprep Yeast Plasmid Miniprep II (Zymo Research Cat. No. D2004).

The resulting DNA was used as template in a 20 µL PCR1 reaction with *RAD54*-specific primers using Q5 High-Fidelity DNA Polymerase (NEB, Cat. No. M0491) to amplify a portion of *RAD54* with universal 5’ adaptor overhangs. Product from PCR1 was used directly as 1:20 template in PCR2 reaction with indexed p5/p7 primers. Thermocycler conditions were as follows for PCR1: 98°C for 30 seconds, 98 °C for 10 seconds, 60 °C for 15 seconds, 72 °C for 15 seconds (steps 2-4 repeated 15 times), 72 °C for two minutes. For PCR2, the annealing temperature was 65 °C and steps 2-4 were repeated 10 times. Barcoded PCR2 reactions were pooled, resolved by 1% agarose gel electrophoresis, and DNA was isolated by Gel Extraction Kit (Qiagen). Paired-end sequencing was performed using an Element AVITI sequencer. Resulting reads were demultiplexed with Bases2Fastq and processed to remove all reads lacking 20-bp sequence upstream and downstream a 9-bp window of interest. This window was extracted, counted, and translated using custom python code. A fold-enrichment score was calculated by comparing the normalized frequency of each amino acid sequence before and after MMS selection. Data was plotted using GraphPad Prism.

### Genetic analysis of Rad54 site-directed mutants

Site-directed mutants of pRS415-*RAD54* were made by assembling 300-bp fragments of *RAD54* (IDT eBlocks) with a PCR amplicon consisting of the remainder of pRS415-*RAD54* (oligos MTP364/365). pRS415-*RAD54* and pRS415-*rad54* with site-directed mutations were transformed into strain yECG46 as described above. After a 72-hour incubation at 30 °C, single colonies were restreaked onto SD –Leu and grown for an additional 72 hours at 30 °C. Cultures inoculated from single colonies were grown overnight in liquid SD – Leu, harvested and resuspended in water to a calculated OD_600_ of 10. Cells were 10-fold serially diluted and 4 µL spotted onto SD –Leu supplemented with MMS as indicated in the figures.

### Genetic analysis of Hed1 point mutants

*HED1* was cloned into the plasmid pYes2 (2μ origin, *URA3* marker, *GAL1* promoter) by generating two fragments by PCR amplification, one from *HED1 S. cerevisiae* W303 genomic DNA with oligos MTP280/MTP281 and another from purified pYes2 with oligos MTP278/MTP279, followed by assembly as above. Site-directed mutants were made by assembling 300-bp fragments of *HED1* (IDT eBlocks) with a PCR amplicon consisting of the remainder of pYes2-*HED1* (oligos MTP366/367).

pYes2-*HED1* and pYes2-*hed1* with site-directed mutations were transformed into strain yECG05 as described above. After a 72-hour incubation at 30 °C, single colonies were restreaked onto SD –Ura and grown for an additional 72 hours at 30 °C. Cultures inoculated from single colonies were grown overnight in liquid SD –Ura and diluted in water to a calculated OD_600_ of 2. Cells were 4-fold serially diluted and 4 µL spotted onto SD –Ura containing either 2% glucose or 2% galactose (Sigma-Aldrich, Cat. No. 91249) supplemented with MMS as indicated in the figures.

### Genetic assays with Rad51 PAP mutants

Wild type *RAD51*, along with an additional 500 base pairs upstream and downstream of the gene, was amplified and cloned into the yeast integrative vector pRS406 utilizing PCR-based methods (primers listed in Table S2). Point mutations were introduced into the *RAD51* gene via site directed mutagenesis and confirmed with long-read sequencing. The vector constructs (including wildtype, mutants, and empty pRS406) were linearized with NcoI-HF restriction enzyme at the *URA3* locus and transformed into yECG48. Integration at the *URA3* locus was confirmed by PCR. Successfully transformed yeast strains were then subjected to sporulation, and the tetrads were dissected. Haploid yeast strains containing a *RAD51* gene at the *URA3* locus, a deletion of *rad51* at the endogenous locus, and of the *MATα* mating phenotype were identified and isolated. These single allele *RAD51* haploids were then used for the spot assays. The spot assays were performed by incubating the strains overnight at 30 °C in liquid YPD media and diluting them to OD_600_ of 1.0 the following day. After additional 10X serial dilutions, the cultures were spotted (4 µL) on freshly poured YPD containing the indicated concentrations of methyl methanesulfonate (MMS; Sigma-Aldrich, Cat. No. 129925) with minimal light exposure. These plates were incubated at 30°C for 2-3 days post spotting and then imaged.

### D-loop assays

D-loop formation experiments were performed in HR buffer (30 mM Tris–OAc [pH 7.5], 50 mM KCl, 20 mM MgOAc, 1 mM DTT, 0.2 mg/ml BSA) using an Atto-647N-labeled tailed DNA duplex consisting of a 21 nt overhang that was homologous for a region on the pUC19 plasmid, and a 56 bp dsDNA region that was not homologous to the plasmid as previously described (Crickard et al., 2020b)(Table S2). Rad51 (WT or triple mutant) (300 nM) was incubated with the tailed duplex DNA (10 nM) in HR buffer supplemented with 10 mM ATP at 30 °C for 15 min. The resulting Rad51-DNA complexes were added to an equal volume of HR buffer containing pUC19 plasmid (9 nM), Rad54 (90 nM), RPA (750 nM) and incubated at 30 °C for 5 min. Reactions were quenched with an equal volume of Stop Buffer (25 mM EDTA, 1% SDS, and 20% glycerol) and then deproteinized by incubation with proteinase K (1 unit) at 37 °C for 20 min. The resulting reaction products were resolved on a 0.9% agarose gel in 1× TAE buffer and detected using a GE Healthcare Life Sciences Typhoon FLA 9500 biomolecular imaging system.

## SUPPLEMENTAL FIGURE LEGENDS

**Figure S1.**
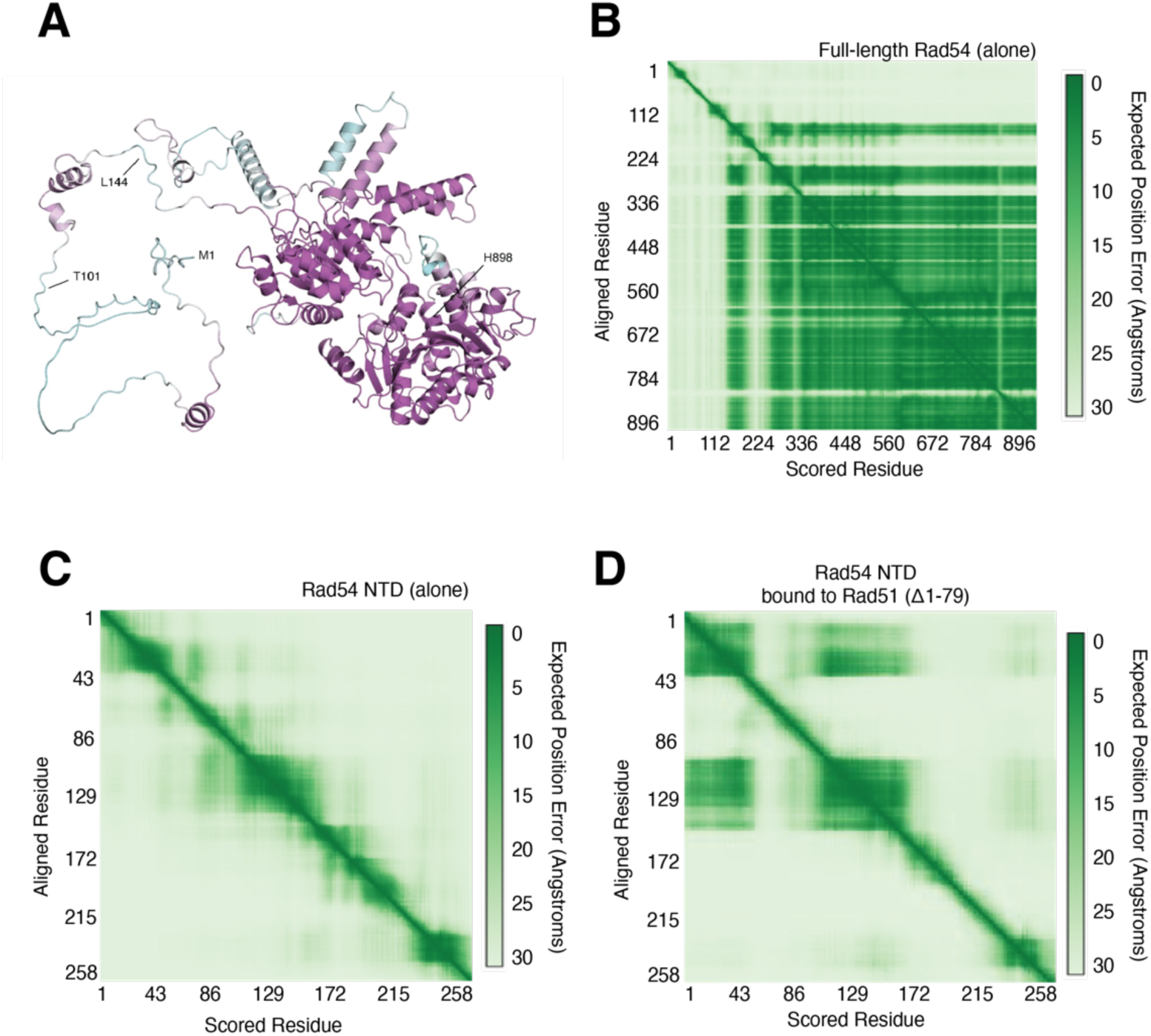
Predicted structure of *S. cerevisiae* Rad54. **(A)** AlphaFold3 predicted structure of full-length *S. cerevisiae* Rad54 alone. **(B)** Predicted aligned error plot for full-length *S. cerevisiae* Rad54 alone. **(C)** Predicted aligned error plot for *S. cerevisiae* Rad54-NTD alone**. (D)** Predicted aligned error plot for *S. cerevisiae* Rad54-NTD bound to the Rad51 dimer. Only the Rad54-NTD region is shown for the comparison to Panel C.

**Figure S2.**
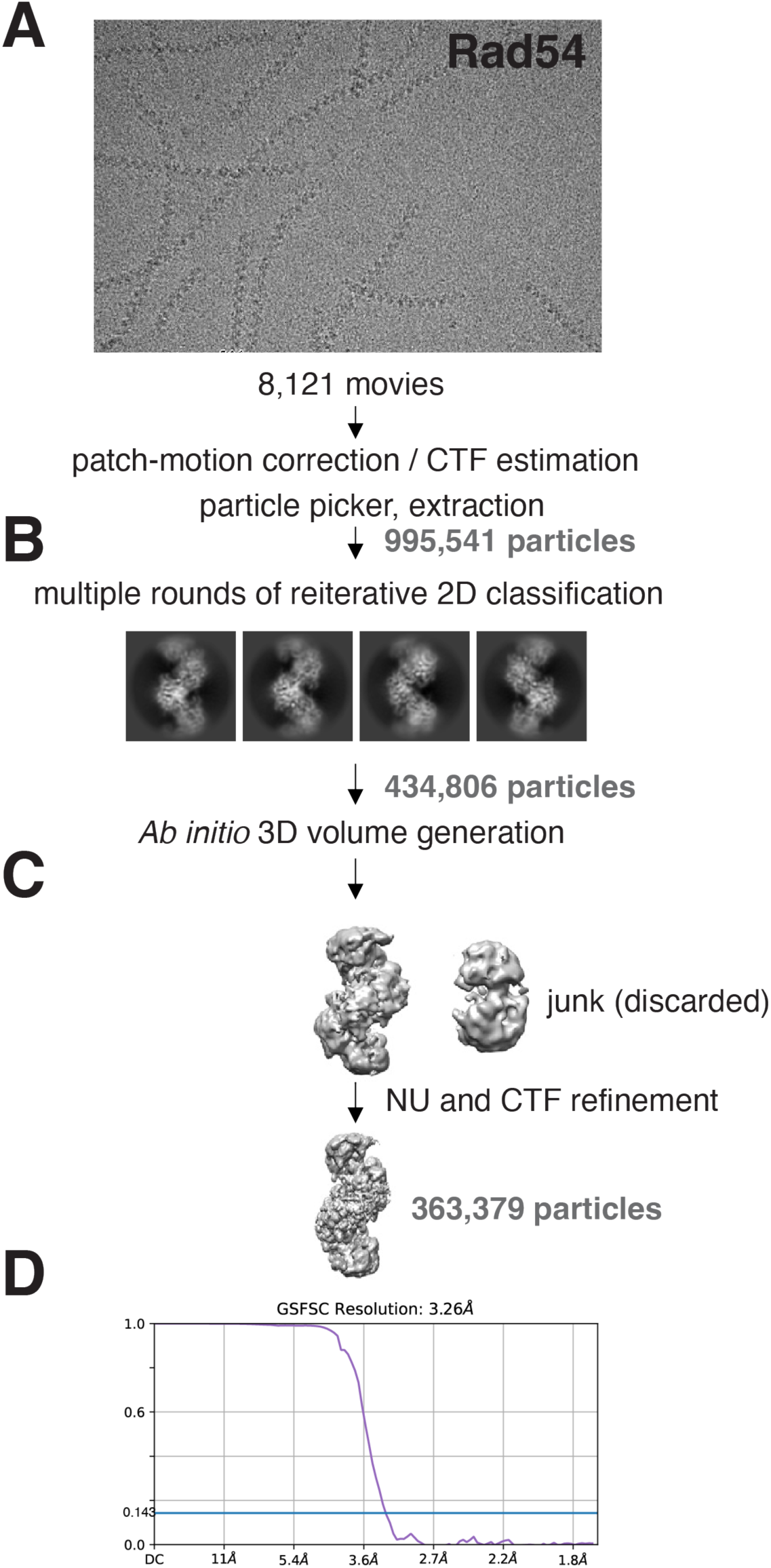
Cryo-EM data processing pipeline for Rad54-Rad51 nucleoprotein filament. **(A)** Representative micrograph used for the Rad51-Rad54 nucleoprotein filament data processing. **(B)** Representative 2D classes selected for 3D classification. **(C)** 3D map generation and refinements. **(D)** Fourier shell correlation curve of the final electron density map.

**Figure S3.**
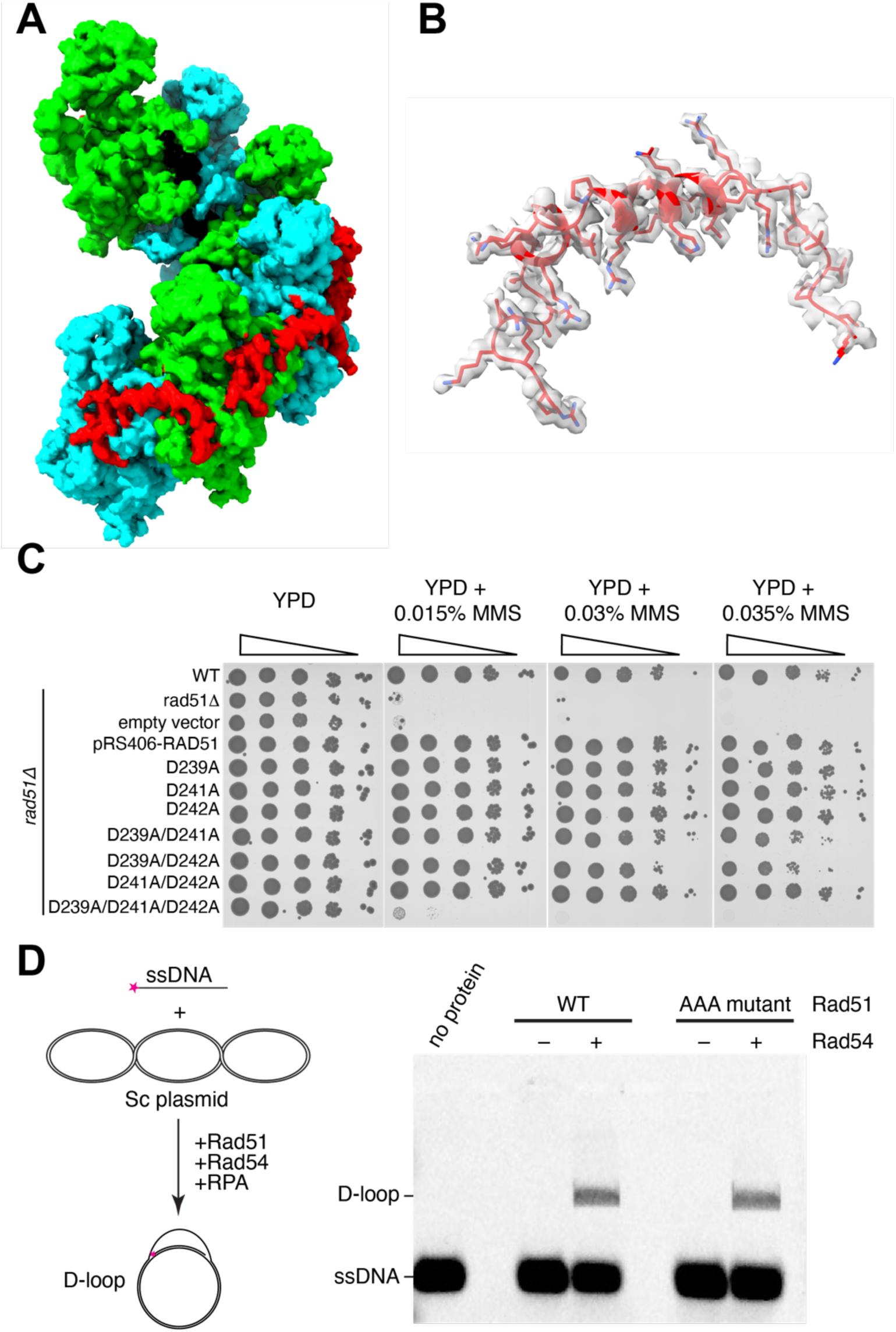
Analysis of the Rad51 interaction domain from Rad54. **(A)** CryoEM density map of the Rad51-Rad54 nucleoprotein filament. Rad51 is shown in alternative cyan and light green, the Rad54 peptide is red, and the ssDNA is shown in black. **(B)** Fitting of the Rad54 peptide into the CryoEM density map. The CryoEM density map of the Rad54 peptide is shown with transparent surface (gray) and the atomic model of the Rad54 peptide is shown as a cartoon and stick model in red. **(C)** Spot assays for single, double, and triple mutants within the Rad51 protruding acidic patch (PAP) motif on media containing the indicated concentrations of MMS. Note, all spot assays were repeated in triplicate. **(D)** Schematic of the *in vitro* D-loop assay using a fluorescently tagged ssDNA substrate and a supercoiled plasmid (left panel) and a D-loop assay showing that the Rad51 PAP triple mutant (D239A, D241A, D242A; denoted as AAA in the figure) retains Rad54-dependent D-loop activity.

**Figure S4.**
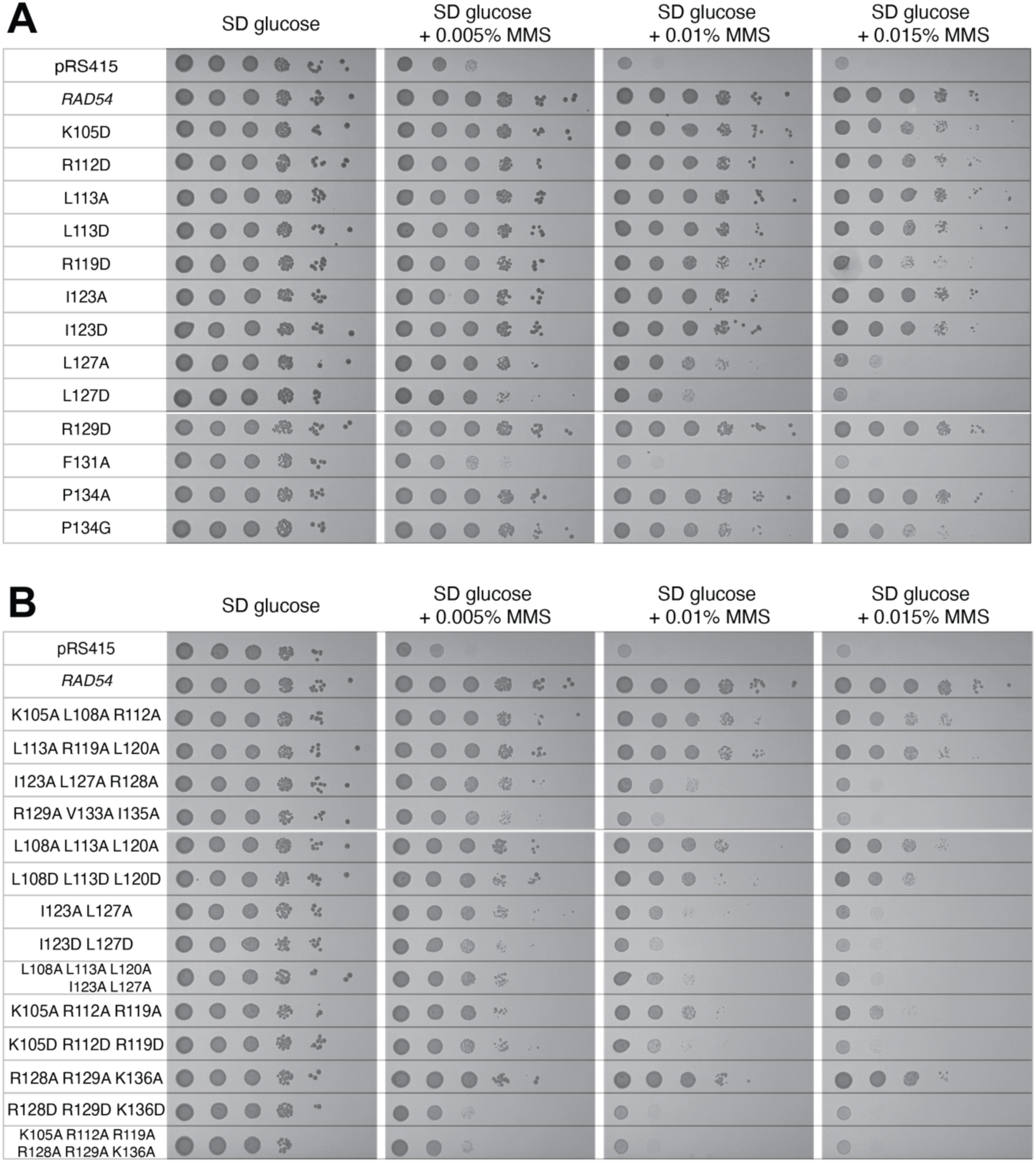
Genetic assays with *rad54* point mutants. **(A)** Spot assays for selected *rad54* single point mutants on media containing the indicated concentrations of MMS. **(B)** Spot assays for *rad54* harboring multiple mutations on media containing the indicated concentrations of MMS. Spot assays are representative of n=3 experiments.

**Figure S5.**
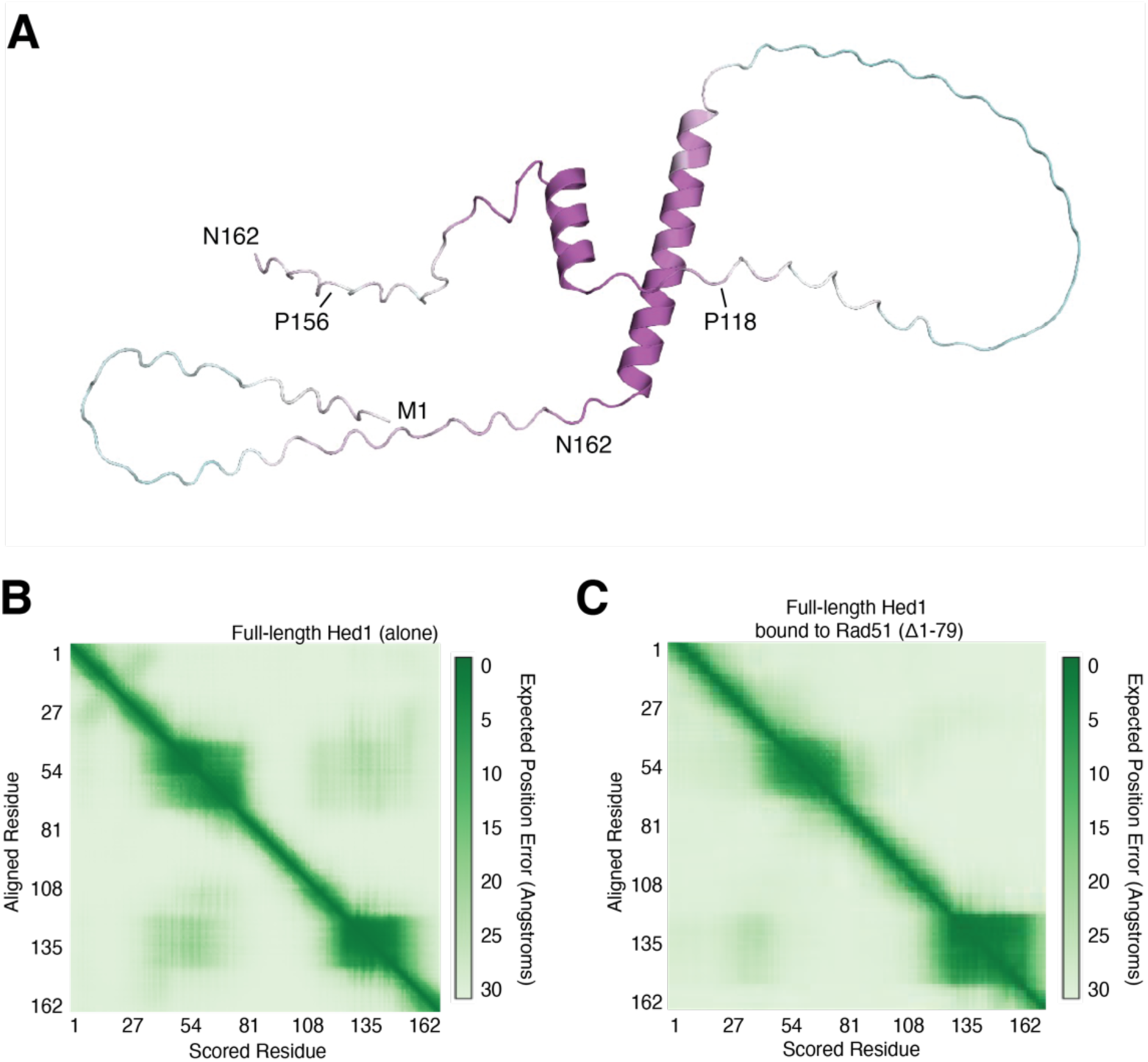
Predicted structure of *S. cerevisiae* Hed1. **(A)** AlphaFold3 predicted structure of full-length *S. cerevisiae* Hed1 alone. **(B)** Predicted aligned error plot for full-length *S. cerevisiae* Hed1 alone. **(C)** Predicted aligned error plot for full-length Hed1 bound to the Rad51 dimer. Only the Hed1 region is shown for the comparison to Panel B.

**Figure S6.**
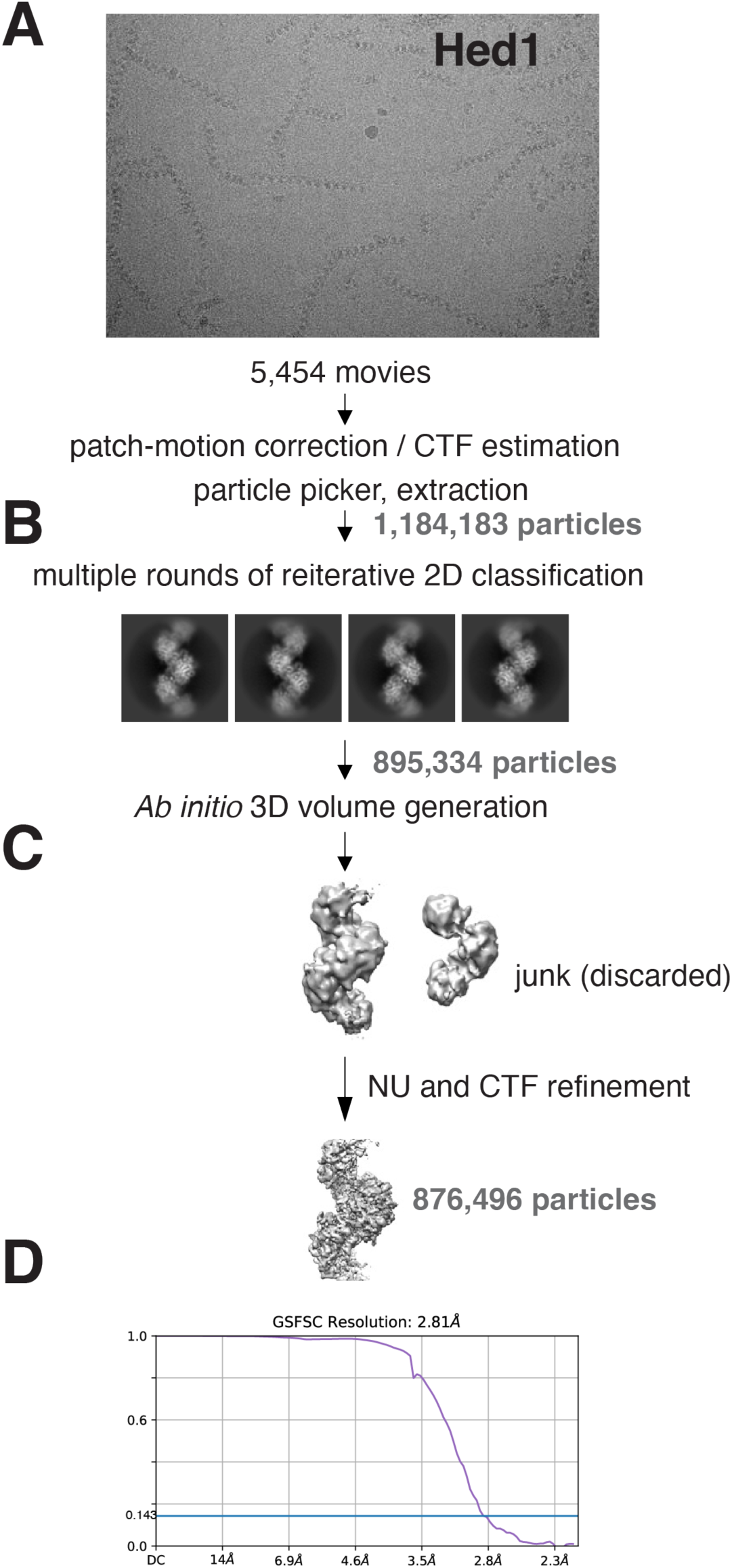
Cryo-EM data processing pipeline for Hed1-Rad51 nucleoprotein filament. **(A)** Representative micrograph used for the Rad51-Hed1 nucleoprotein filament data processing. **(B)** Representative 2D classes selected for 3D classification. **(C)** 3D map generation and refinements. **(D)** Fourier shell correlation curve of the final electron density map.

**Figure S7.**
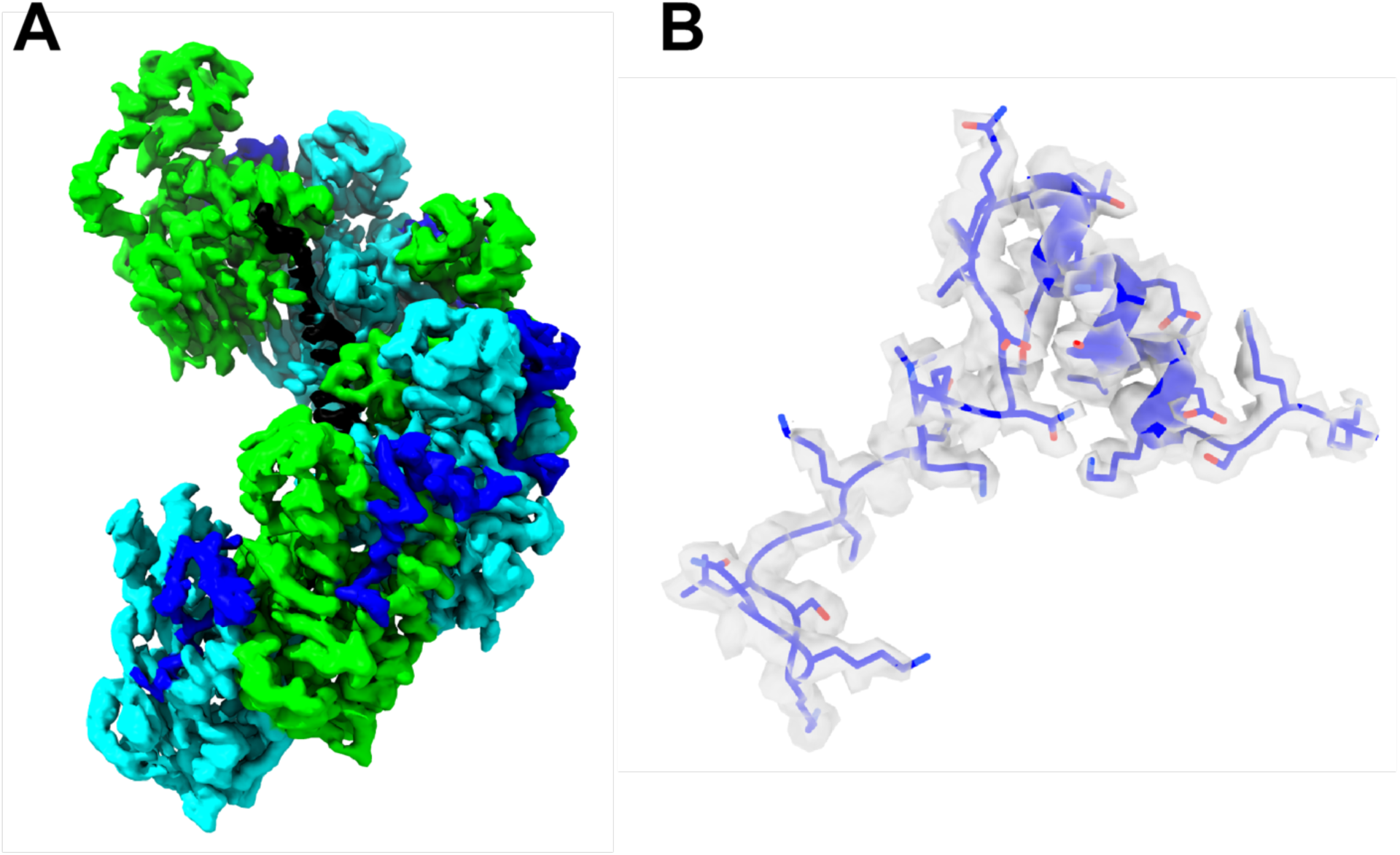
Electron density for the Rad51 interaction domain from Hed1. **(A)** CryoEM density map of the Rad51-Hed1 nucleoprotein filament. Rad51 is shown in alternating cyan and light green, the Hed1 peptide is dark blue, and the ssDNA is shown in black. **(B)** Fitting of the Hed1 peptide into the CryoEM density map. The CryoEM density map of the Rad54 peptide is shown with transparent surface (gray) and the atomic model of the Rad54 peptide is shown as a cartoon and stick model in blue.

**Figure S8.**
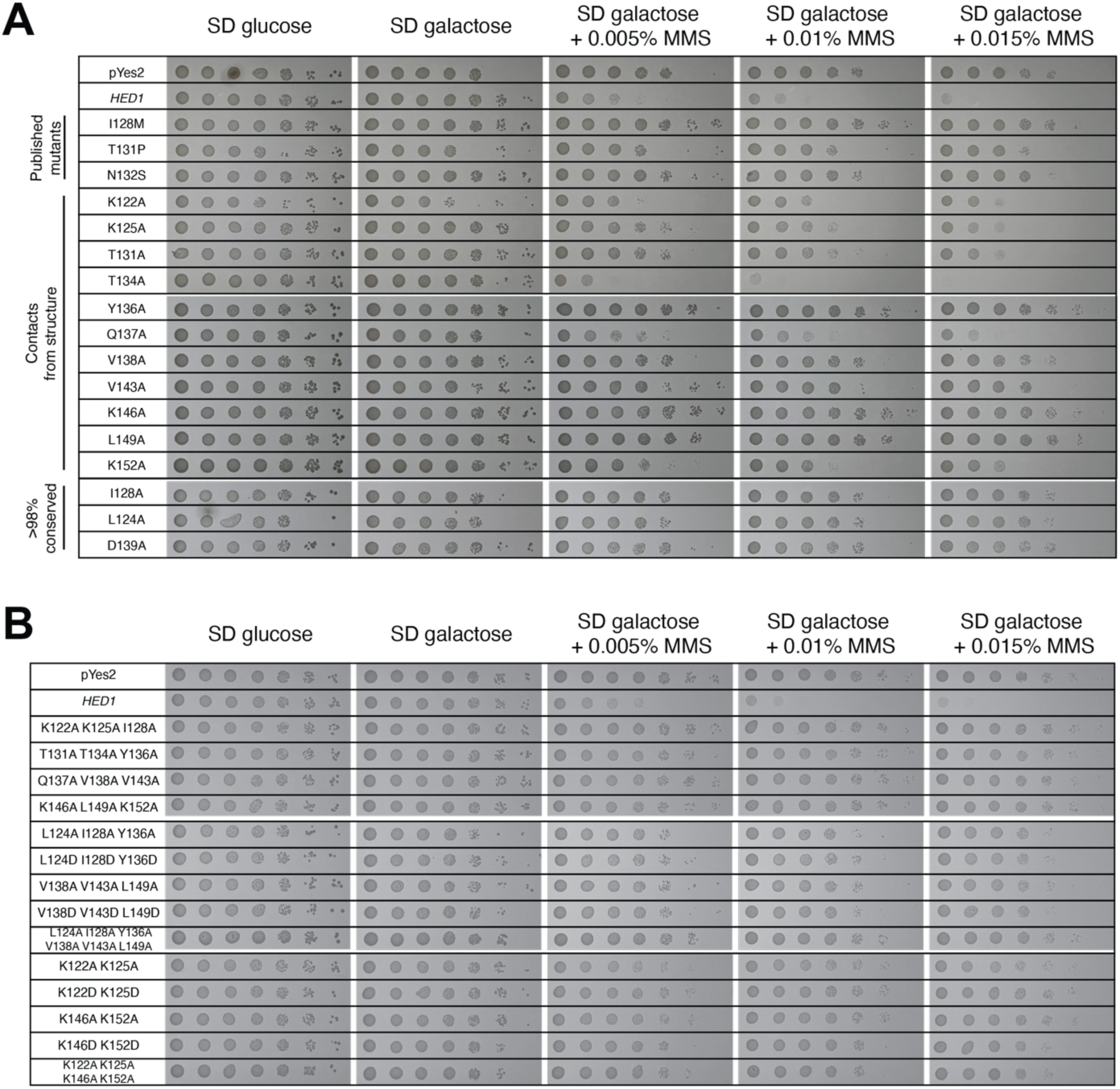
Genetic assays with *hed1* point mutants. **(A)** Spot assays for selected *hed1* single point mutants on media containing the indicated concentrations of MMS. **(B)** Spot assays for *hed1* harboring multiple mutations on media containing the indicated concentrations of MMS. In these assays, Hed1 expression was suppressed in the presence of glucose, whereas Hed1 expression was induced by the inclusion of galactose in the growth media. Spot assays are representative of n=3 experiments.

**Figure S9.**
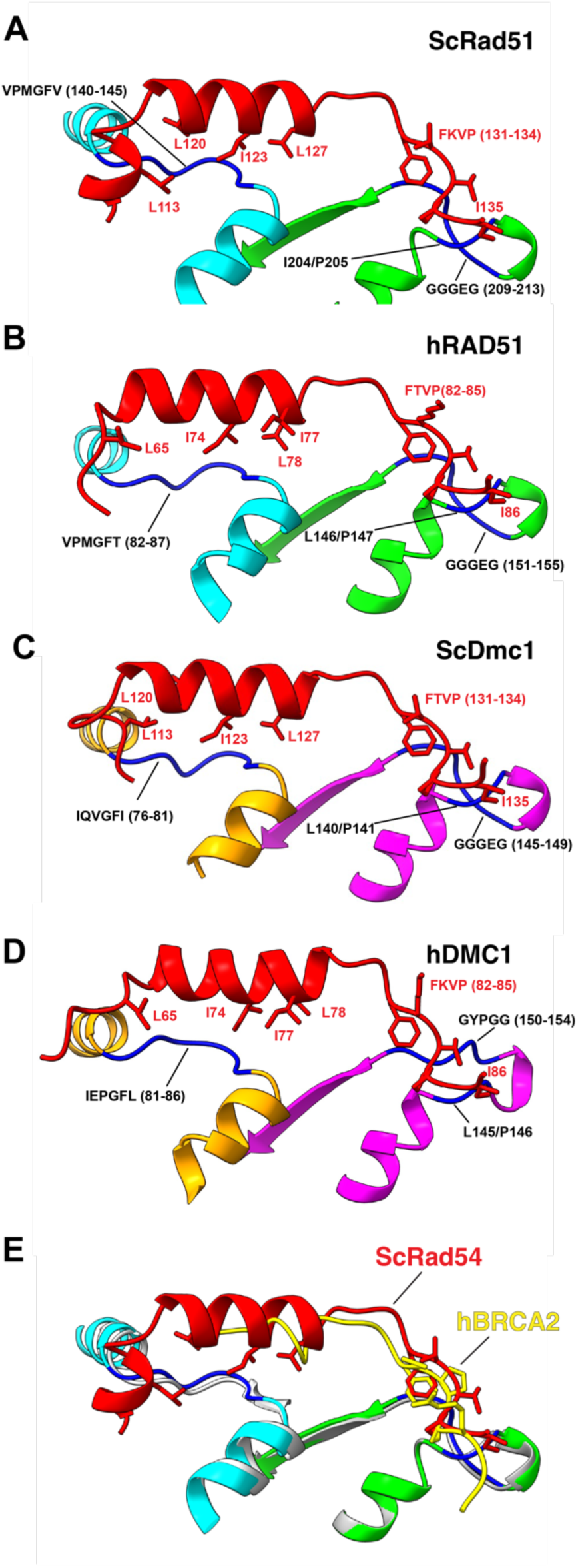
Conservation of the Rad54 interaction with different eukaryotic recombinases. **(A)** CryoEM structure of the *S. cerevisiae* Rad51 interaction motif from Rad54 (in red) bound to two adjacent Rad51 monomers (cyan and green). **(B)** AlphaFold3 prediction of the Rad51 interaction motif from human RAD54 bound to two adjacent human RAD51 monomers (cyan and green). **(C)** AlphaFold3 prediction of the Rad51 interaction motif from *S. cerevisiae* Rad54 bound to two adjacent *S. cerevisiae* Dmc1 monomers (orange and magenta). **(D)** AlphaFold3 prediction of the Rad51 interaction motif from human RAD54 bound to two adjacent human DMC1 monomers (orange and magenta). Conserved regions of contact are indicated in all panels. **(E)** Overlay of the CryoEM structure for *S. cerevisiae* Rad51 interaction motif from Rad54 (in red) bound to two adjacent Rad51 monomers (cyan and green) with human RAD51 (in gray) bound to the TR2 motif from BRCA2 (in yellow; PDB accession codes 8PBC)(Appleby et al., 2023).

**Table S1.**

|  | **Rad51_Rad54**<br>**PDB: 9E6L**<br>**(EMD-47572)** | **Rad51_Hed1**<br>**PDB: 9E6N**<br>**(EMD-47573)** |
| --- | --- | --- |
| **Data collection and processing** |  |  |
| Microscope | Titan Krios | Titan Krios |
| Voltage (keV) | 300 | 300 |
| Detector | K3 | K3 |
| Magnification | 105,000 | 85,000 |
| Voltage (kV) | 300 | 300 |
| Electron exposure (e<sup>-</sup>/Å<sup>2</sup>) | 59.51 | 51.21 |
| Defocus range (μm) | -0.8 to -2.5 | -0.8 to -2.5 |
| Pixel size (Å) | 0.844 | 1.083 |
| Initial particles picked | 995,541 | 1,184,183 |
| Final particles used | 363,379 | 876,496 |
| Map resolution (Å) | 3.3 | 2.8 |
| FSC threshold | 0.143 | 0.143 |
| Map resolution range (Å) | 3.3-3.7 | 2.8-3.3 |
| **Refinement** |  |  |
| Model resolution (Å) | 3.3 | 2.8 |
| FSC threshold | 0.143 | 0.143 |
| *Model composition* |  |  |
| Non-hydrogen atoms | 16,562 | 16,759 |
| Protein residues | 2,092 | 2,115 |
| Ligands | ATP, MG | ATP, MG |
| *R.m.s. deviations* |  |  |
| Bond lengths (Å) | 0.002 | 0.003 |
| Bond angles (°) | 0.453 | 0.467 |
| *Validation* |  |  |
| MolProbity score | 1.27 | 1.25 |
| Clash score | 5.14 | 4.81 |
| Rotamer outliers (%) | 0 | 0 |
| *Ramachandran plot* |  |  |
| Favored (%) | 98.69 | 98.42 |
| Allowed (%) | 1.26 | 1.58 |
| Outliers (%) | 0 | 0 |

## Notes

### Competing Interest Statement

The authors have declared no competing interest.

